# Phylogeny and Multiple Independent Whole-Genome Duplication Events in the Brassicales

**DOI:** 10.1101/789040

**Authors:** Makenzie E. Mabry, Julia M. Brose, Paul D. Blischak, Brittany Sutherland, Wade T. Dismukes, Christopher A. Bottoms, Patrick P. Edger, Jacob D. Washburn, Hong An, Jocelyn C. Hall, Michael R. McKain, Ihsan Al-Shehbaz, Michael S. Barker, M. Eric Schranz, Gavin C. Conant, J. Chris Pires

## Abstract

Whole-genome duplications (WGDs) are prevalent throughout the evolutionary history of plants. For example, dozens of WGDs have been phylogenetically localized across the order Brassicales, specifically, within the family Brassicaceae. However, while its sister family, Cleomaceae, has also been characterized by a WGD, its placement, as well as that of other WGD events in other families in the order, remains unclear. Using phylo-transcriptomics from 74 taxa and genome survey sequencing for 66 of those taxa, we infer nuclear and chloroplast phylogenies to assess relationships among the major families of the Brassicales and within the Brassicaceae. We then use multiple methods of WGD inference to assess placement of WGD events. We not only present well-supported chloroplast and nuclear phylogenies for the Brassicales, but we also putatively place Th-α and provide evidence for previously unknown events, including one shared by at least two members of the Resedaceae, which we name Rs-α. Given its economic importance and many genomic resources, the Brassicales are an ideal group to continue assessing WGD inference methods. We add to the current conversation on WGD inference difficulties, by demonstrating that sampling is especially important for WGD identification.

## INTRODUCTION

The Brassicales are an economically important order of flowering plants, home to many crop species such as kale, broccoli, cabbage, cauliflower, papaya, capers, and canola as well as several model plants including *Arabidopsis*. Currently, there are 17 accepted families within the Brassicales (APG IV 2016), with the family Brassicaceae receiving most attention due to the many crop species and model plants placed within it. The two most closely related families to Brassicaceae, Cleomaceae and Capparaceae, have received much less attention: however, collectively these three families comprise 94% of the order (Edger et al. 2015). Sister to these three families is a polytomy of four families; Tovariaceae, Gyrostemonaceae, Resedaceae, and Pentadiplandraceae. This clade is then followed by Emblingiaceae, [[Salvadoraceae + Bataceae], Koeberliniaceae], Limnanthaceae, Setchellanthaceae, [Caricaceae + Morginaceae], and [Tropaeolaceae + Akaniaceaeae] (**Supp. Figure 1**; APG IV 2016). Together, the order is dated around 103 mya, with extant species contributing to 2.2% of the total core eudicot diversity (Magallon et al. 1999; Cardinal-McTeague et al. 2016). Previous research has identified multiple whole-genome duplication (WGD) events across the order using a variety of comparative methods, including genomics, transcriptomics, and molecular cytogenetics (Vision et al. 2000; Schranz & Mitchell-Olds 2006; Barker et al. 2009; Cheng et al. 2013; Kagale et al. 2014; Edger et al. 2015; Edger et al. 2018; Lysak 2018). Four of the most studied events include one near the base of the order (At-β; Edger et al. 2015; Edger et al. 2018), at the base of the Brassicaceae family (At-α; Vision et al. 2000; Haudry et al. 2013; Edger et al. 2015), a triplication event the base of the tribe Brassiceae in the Brassiaceae (Lysak et al. 2005), and an unplaced event within the Cleomaceae (Th-α; Schranz & Mitchell-Olds 2006; Barker et al. 2009).

The Brassicaceae family has the largest number of accepted species with >4,000 named (BrassiBase). It contains the model plant organism, *Arabidopsis thaliana* (Arabidopsis Genome Initiative 2000), as well as the important crops of the *Brassica* and *Raphanus* groups. The clades of this family have been placed into three major lineages (Lineage I, Lineage II, and Lineage III; Beilstein et al. 2006), with notable named clades acknowledged more recently (Huang et al. 2016; Nikolov et al. 2019). However, the relationships among these lineages and clades are still unclear. Besides work in understanding the relationships within the Brassicaceae, another major area of research within the family has been on the considerable glucosinolate diversity (Kliebenstein et al. 2001, Ratzka et al. 2002, Züst et al. 2018). Many aspects of these plant defense compounds have been studied, including their coevolution with insect herbivores (Edger et al. 2015). Yet another major area of research in the family is on investigating the impact of WGDs events. The Brassicaceae seems to be especially enriched with WGD events both at the base and within the family (Barker et al. 2009; Kagale et al. 2014; Edger et al. 2015; Mandakova et al. 2017; Edger et al. 2018).

Sister to the Brassicaceae is the Cleomaceae, an herbaceous family of ~270 species of pantropical plants that diverged from the Brassicaceae around 40 mya (Edger et al. 2015). The Cleomaceae display a much wider range of floral morphologies than its sister family, a fact which has been the focus of several studies (Bhide et al. 2014; Brock 2014; Bayat et al. 2018). This family is unique in the Brassicales as it contains species with C4 photosynthesis (*Gynandropsis gynandra* and *Coalisina angustifolia*, formally *Cleome angustifolia*) as well as, though not unique to Cleomaceae (Schlüter et al. 2016), a C3-C4 intermediate (*Coalisina paradoxa*, formally *C. paradoxa*; van den Bergh et al. 2014). The Cleomaceae are known to have at least one independent polyploidy event that occurred after their split from the Brassicaceae, named Th-α after *Tarenaya hassleriana.* It has been dated to around 13.7 mya (Schranz & Mitchell-Olds 2006; Barker et al. 2009; Cheng et al. 2013). We note, however, that the analyses used for this identification and dating used only partial genomic fragments, ESTs, or a single genome. Van den Bergh et al. (2014) later determined that this event was shared with at least the species *G. gynandra*, a C_4_ species, and, more recently, it was determined that Th-α was not shared with *Cleome violacea* (Emery et al. 2018). Although this duplication has been identified and is shared with at least two members of the Cleomaceae, it remains a mystery as to where the Th-α event occured within the context of the phylogeny for the family (van den Bergh et al. 2014; Bayat et al. 2018).

The sister family to the Brassicaceae and Cleomaceae, the Capparaceae -a mostly woody tropical family of 450 species, is much less studied than the two families just discussed. Like the Cleomaceae, they are also very diverse in their floral morphology (Endress 1992). The Capparaceae produce glucosinolates (as do all members of the order), however, they share the production of unique methyl-glucosinolates with only the Cleomaceae (Hall et al. 2002; Mithen et al. 2010). In this group of economic importance is the plant species *Capparis spinosa*, or capers. Recent work using chromosome counts hypothesized that the Capparaceae and a more distant family, the Resedaceae, may too possess unique WGD events (Lysak 2018). The Resedaceae, a relatively small clade of ~ 85 species, are mostly distributed across Europe, the Middle East, and Africa with one taxon occurring in North America (*Oligomeris linifolia*) due to a long-distance dispersal event (Martin-Bravo et al. 2007; 2009; Cardinal-McTeague et al. 2016).

To infer phylogenetic relationships within the Brassicales, we use phylo-transcriptomics, a quickly evolving subdiscipline of phylogenomics that uses RNA-seq data (Dunn et al. 2008; McKain et al. 2012; Yang et al. 2015; Washburn et al. 2017). Transcriptomics allows access to many more nuclear genes than using traditional PCR but is less expensive than sequencing an entire genome. Using RNA-seq data not only allows for the inference of phylogenetic relationships, but also allows for assessing gene and genome duplication events (Baker et al. 2009; McKain et al. 2012). However, one major difficulty in using transcriptomes for phylogenetic inference is the problem of determining orthology. To address this, several methods have been developed, including those that aim to identify orthogroups, or sets of genes that are descended from a single gene in the last common ancestor of the group or species of interest (Duarte et al. 2010, Emms and Kelly 2015). Here we use OrthoFinder2 (Emms and Kelly 2018), because it offers both improvements in orthogroup inference accuracy, and also in computational speed, especially when using Diamond (Buchfink et al. 2015). Methods like these have helped enable phylo-transcriptomics to be extremely useful for inferring species relationships, understanding gene evolution, and elucidating WGD events.

With WGD events well established across the Brassicaceae family, including At-α at the base (Vision et al. 2000; Edger et al. 2015), and the identification of a unique and more recent, yet unplaced event in the Cleomaceae (Th-α; Schranz & Mitchell-Olds 2006; Barker et al. 2009; Cheng et al. 2013), the Brassicales are an intriguing group for the study of polyploidy. Using phylo-transcriptomics with a focus on sampling the Brassicaceae and Cleomaceae, with additional sampling of the Capparaceae, Resedaceae, Bataceae, Caricaceae, and Moringaceae families, we aim to answer remaining questions on the placement of events including Th-α. In particular, we ask if Th-α is shared across the Cleomaceae, or if this family, like the Brassicaceae, is characterized by multiple events. We also test the recent hypothesis that the families Resedaceae and Capparaceae possess independent WGD events (Lysak 2018). Together, it is clear that the Brassicales are a powerful resource for the study of WGD, and will be an important group to further test how WGD correlates with traits of interest, such as variation in floral morphology, photosynthesis types, or metabolism.

## RESULTS

### Sequence Matrices

DNA read pools ranged in size from 6,637,717 to 13,335,392 reads. We also analyzed two previously sequenced copies of matK and ndhF in combination with our own data, resulting in alignment lengths of 1,521 and 985 bp for each gene, respectively. After assembly of complete chloroplasts, the inferred genomes for 66 taxa ranged in length from 137,110 to 160,272 bp. The large single copy (LSC), small single copy (SSC), and inverted repeat (IR) regions were isolated and aligned separately, with total alignment lengths of 84,350 bp, 17,931 bp, and 26,500 bp, respectively. Both chloroplast analyses had 100% occupancy for taxa included.

RNA read pools ranged in size from 5,555,024 to 59,723,745 reads, with an average of 22,520,865 reads. To check completeness of transcriptomes, assemblies were run though BUSCO v3 (Simão et al. 2015; Waterhouse et al. 2017). All assemblies had greater than 66% complete genes with less than 12% of genes missing or fragmented (**Supp. Figure 2**). Using OrthoFinder v.2.2.6 (Emms & Kelly 2018) we recovered 47,600 orthogroups across the Brassicales. Filtering for an 80% taxon occupancy (59/74 taxa) yielded 10,968 orthogroups. After filtering for alignment quality by allowing for only 40% gaps, we recovered 2,663 orthogroups. Finally, pruning trees for any remaining paralogs by using a minimum of 10 taxa as a cutoff resulted in 1,284 orthogroups which were then used for species tree inference. Following the steps above for each family (Brassicaceae, Capparaceae, Cleomaceae, and a group of Resedaceae + Bataceae + Moringaceae + Caricaceae) we recovered 2,100, 10,214, 3,626, and 8,476 orthogroups, respectively (**Supp. Table 1)**.

### Phylogenomics of the Brassicales

In the analysis of just two chloroplast genes, matK and ndhF, of 91 taxa from the study by Hall (2008) and all 66 of our samples we recover the same overall relationships as published for other chloroplast phylogenies of the Brassicales (Hall 2008; Cardinal-McTeague et al. 2016; Edger et al. 2018; **Supp. Figure 3**). Overall, this tree is recovered with many poorly supported nodes and importantly, for a few species that were included in both this and the Hall (2008) study, placement in the tree is paraphyletic. This includes the species *Stanleya pinnata*, *Cleomella lutea*, *Andinocleome pilosa*, and *Capparis tomentosa*. This lack of congruence for species placement may be due to the fact that species are mislabeled (*Cleomella lutea*), poor species descriptions, or species being more genetically diverse than previously thought. Due to this uncertainty in taxon identification, we refer to these samples as *Brassicaceae sp*, *Polanisia sp.*, *Cleomaceae sp*, and *Capparaceae sp*, respectively.

For whole-chloroplast analyses, using just one copy of the IR, all nodes except four are recovered with 70% bootstrap support or better with a topology which is largely congruent with previous studies (Hall 2008; Cardinal-McTeague et al. 2016; Edger et al. 2018). This includes a clade of *Moringa oleifera* and *Carica papaya* sister to a clade of [Bataceae + Resedaceae + Capparaceae + Cleomaceae + Brassicaceae], followed by Bataceae sister to [Resedaceae + Capparaceae + Cleomaceae + Brassicaceae], Resedaceae sister to [Capparaceae + Cleomaceae + Brassicaceae], and finally, Capparaceae sister to [Cleomaceae + Brassicaceae](**Supp. Figure 4**). Relationships among the major lineages within the Brassicaceae are also in agreement with previous studies (Guo et al. 2017). We recover *Aethionema arabicum* as sister to the rest of the family, followed by Lineage I sister to [Lineage III + Clade C + Lineage II and Expanded Lineage II] and Lineage III sister to [Clade C + Lineage II and expanded Lineage II]. Within the Cleomaceae relationships are mostly congruent with previous studies (Hall 2008; Patchell et al. 2014), with the exception of the placement of *Polanisia* sister to *Cleome* sensu stricto (after Patchell et al. 2014) rather than the rest of the family. Most likely due to sampling, our relationships among the Capparaceae are not congruent with previous studies (Hall 2008, Tamboil et al. 2018). Previous studies with more sampling recover *Boscia sp.* sister to *Cadaba*, while in our study we recover *Boscia* sister to *Capparis*.

Analysis of nuclear data from the transcriptome with ASTRAL-III recovered a well-resolved tree with all nodes but four recovered with a local posterior probability of 0.7 or higher (Figure 1). The overall relationships of the families and major lineages are congruent with previous studies using transcriptomics (Edger et al. 2015). As with the whole-chloroplast phylogeny, we recover a clade of *Moringa oleifera* and *Carica papaya* sister to a clade of [Bataceae + Resedaceae + Capparaceae + Cleomaceae + Brassicaceae], Bataceae sister to [Resedaceae + Capparaceae + Cleomaceae + Brassicaceae], Resedaceae sister to [Capparaceae + Cleomaceae + Brassicaceae], and finally, Capparaceae sister to [Cleomaceae + Brassicaceae]. Within Brassicaceae, the major lineages are recovered as supported by previous literature (Huang et al. 2016; Nikolov et al. 2019) with *Aethionema arabicum* as sister to the rest of the family, followed by Lineage III sister to [Lineage I + Clade C + Lineage II and expanded Lineage II] and Lineage I sister to [Clade C + Lineage II and expanded Lineage II]. Within Cleomaceae, the relationships were also mostly congruent with previous nuclear phylogenies (Patchell et al. 2014; Cardinal-McTeague et al. 2016) with the same exception of the placement of *Polanisia* in relation to the other clades of Cleomaceae as discussed above. Finally, our limited sampling of the Capparaceae again limits our ability to say much about the relationships within the family, however to date there is no phylogeny for the family based solely on nuclear data.

### Known WGD Events Not Recovered with Strong Support When Sampling Across the Brassicales

Two of the most popular methods used to detect WGD include phylogenomics - using individual gene tree topologies, gene counts, and a known species tree - and Ks plots, which allow for the identification of signatures left behind in paralogs after WGD. In order to provide multiple lines of evidence for novel WGD events. We use a combination of these two approaches to test hypotheses of proposed WGD across the Brassicales. Using PUG (github.com/mrmckain/PUG), a phylogenomic WGD estimation method, resulted in the recovery of some known events with high support (e.g. At-α and At-β), yet failed to produce strong support for other known events such as the Brassiceae triplication event when including all taxa (Figure 1). We note that PUG does indicate that there are 65 unique gene duplications that match that node when considering gene trees with 80% bootstrap support. However, when compared to other known events (At-α and At-β) with counts over 300 and 150 respectively, this is surprisingly low. Therefore, to increase the number of orthogroups used to infer species trees, as well as increase the number of gene trees to query putative paralogs against, we further broke down analyses to the familial level. By running the Brassicaceae, Capparaceae, Cleomaceae, and [Resedaceae + Bataceae + Moriagaceae + Caricaceae] families separately, we are able to improve WGD detection of known events.

**Figure 1.**
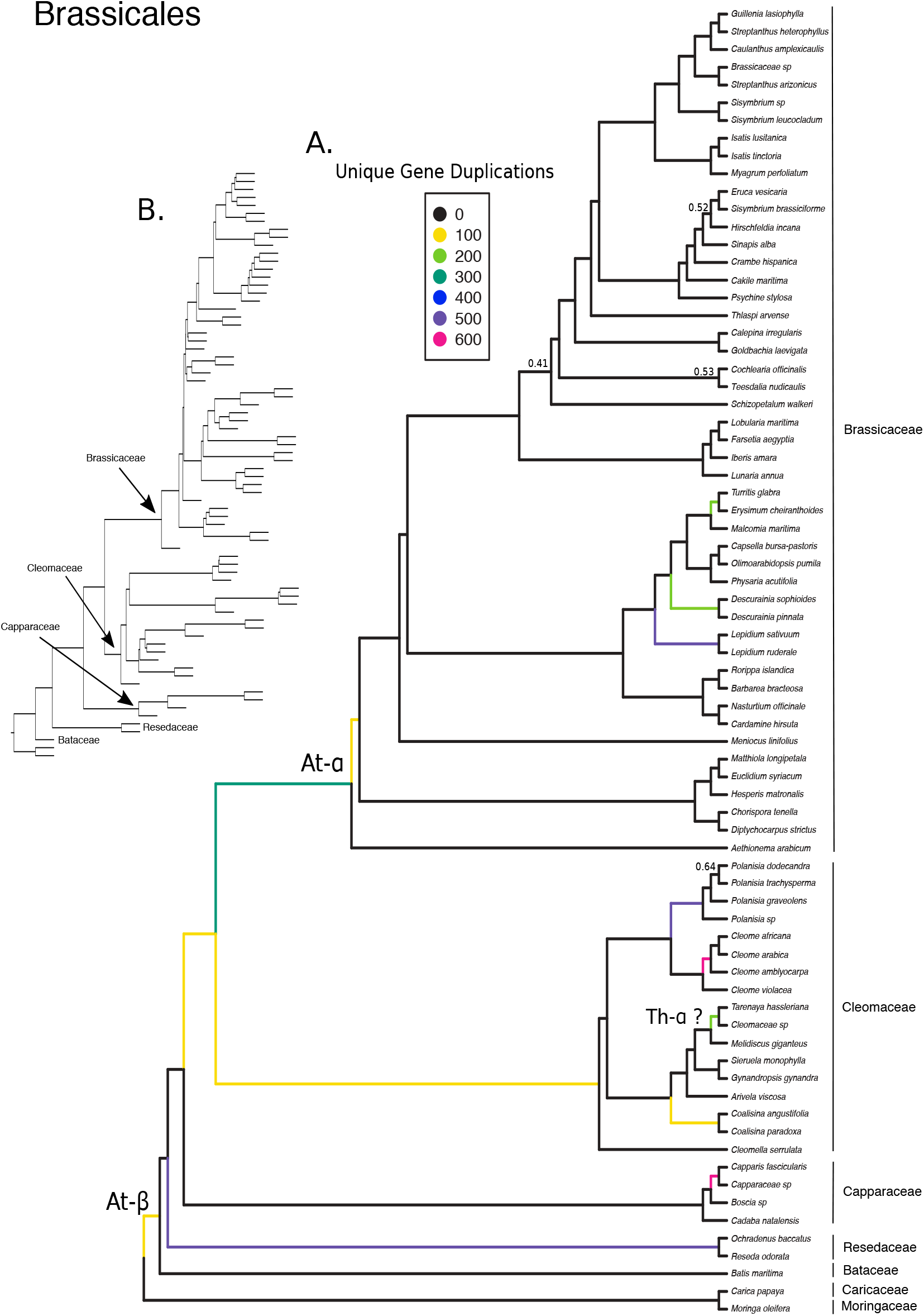
Coalescent-based species phylogeny and whole-genome duplication events of the Brassicales. **(A)** Coalescent-based species tree with branch lengths proportional. Known events (At-α and At-β) are indicated, as well as possible placement of Th-α. Branches colored by number of unique gene duplications as determined by PUG (github.com/mrmckain/PUG). Support values are indicated if below 0.7 local posterior probability. **(B)** Coalescent-based species tree with branch lengths.

### Recovery of Known WGD Events in the Brassicaceae

Analysis of just the Brassicaceae family identifies not only At-α at the base of the family, but also successfully identifies the Brassiceae triplication event (Lysak et al. 2005; Figure 2). We also recover up to five additional neopolyploid events between; 1) *Chorispora tenella* and *Diptychocarpus strictus*, 2) *Lepidium ruderale* and *L. sativum*, 3) *Descurainia sophioides* and *D. pinnata*, 4) *Turritis glabra* and *Erysimum cheiranthoides*, and 5) a clade of *Isatis lusitanica*, *I. tinctoria*, and *Myagrum perfoliatum*.

**Figure 2.**
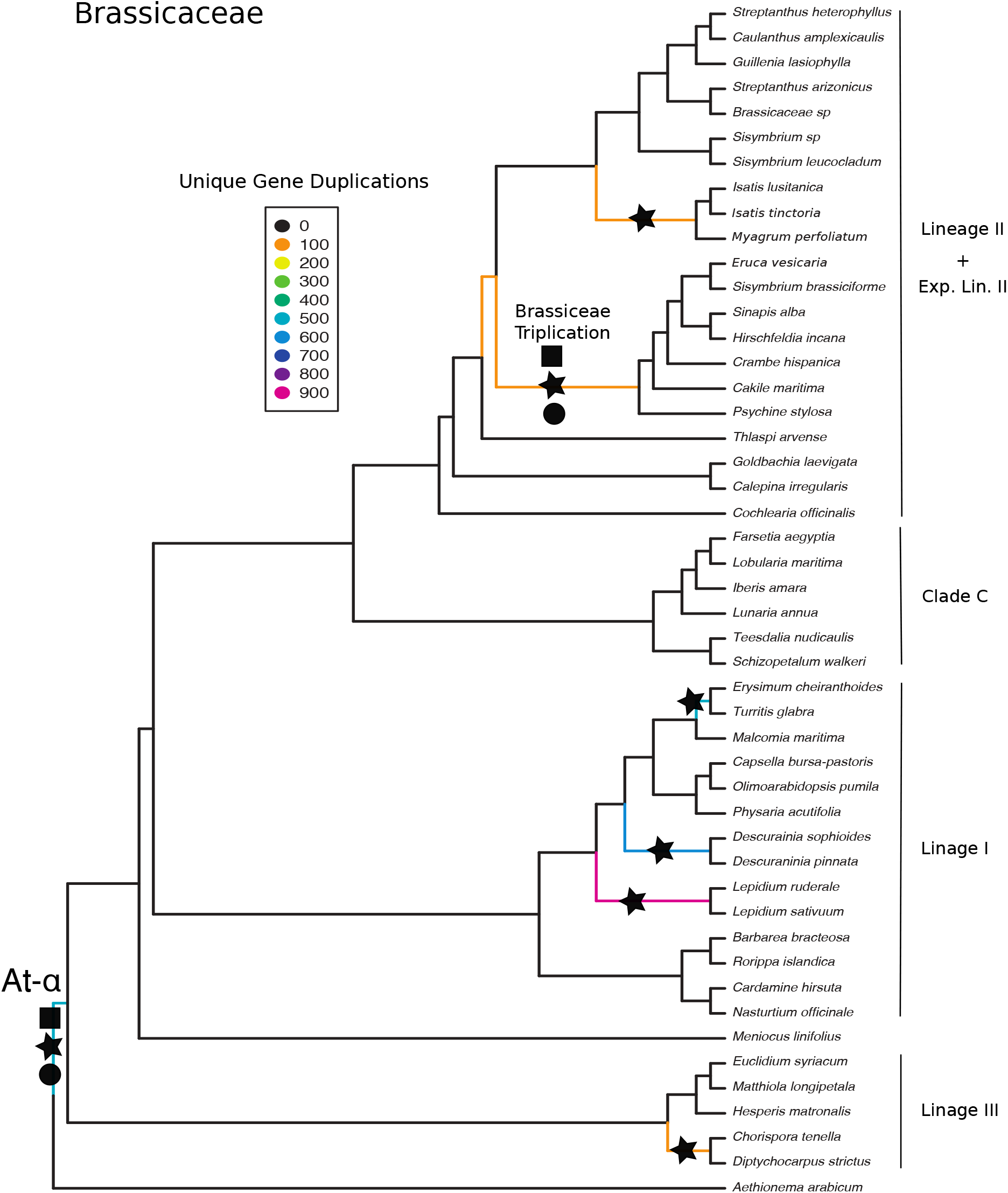
Coalescent-based species phylogeny and whole-genome duplication events of the Brassicaceae. Branches colored by number of unique gene duplications as determined by PUG (github.com/mrmckain/PUG). Black stars indicate WGD events identified by PUG, Black square indicate WGD events identified by FASTKs (McKain et al. 2016), and Black circles indicate WGD events identified by DupPipe (Barker et al. 2010). Support values are all above 0.7 local posterior probabilities.

Ks plots, run using both FASTKs to estimate pairwise Ks values (github.com/mrmckain/FASTKs; McKain et al. 2016) and DupPipe to estimate Ks values using duplications in gene trees (Barker et al. 2010), mostly show agreement with the WGD events inferred by the phylogenetic method, PUG. For example, within the Brassicaceae, Ks plots from both analyses recover the Brassiceae Triplication (Ks ~ 0.3; **Supp. Figure 5**). However, for the neopolyploid events within the Brassicaceae and At-α, Ks plots show differing results between FASTKs and DupPipe, some with and others without evidence for WGD events (**Supp. Figure 5**).

### Difficulty in Placement of Th-α in Cleomaceae

When running PUG using just the Cleomaceae family members, we place Th-α as potentially shared between *Tarenaya hassleriana* and *Cleomaceae sp*. We also identify up to three additional events between; 1) *Coalisina paradoxa and Coalisina angustifolia*, 2) four species of *Polanisia*, and 3) *Cleome amblyocarpa*, *Cleome africana*, and *Cleome arabica* (Figure 3).

**Figure 3.**
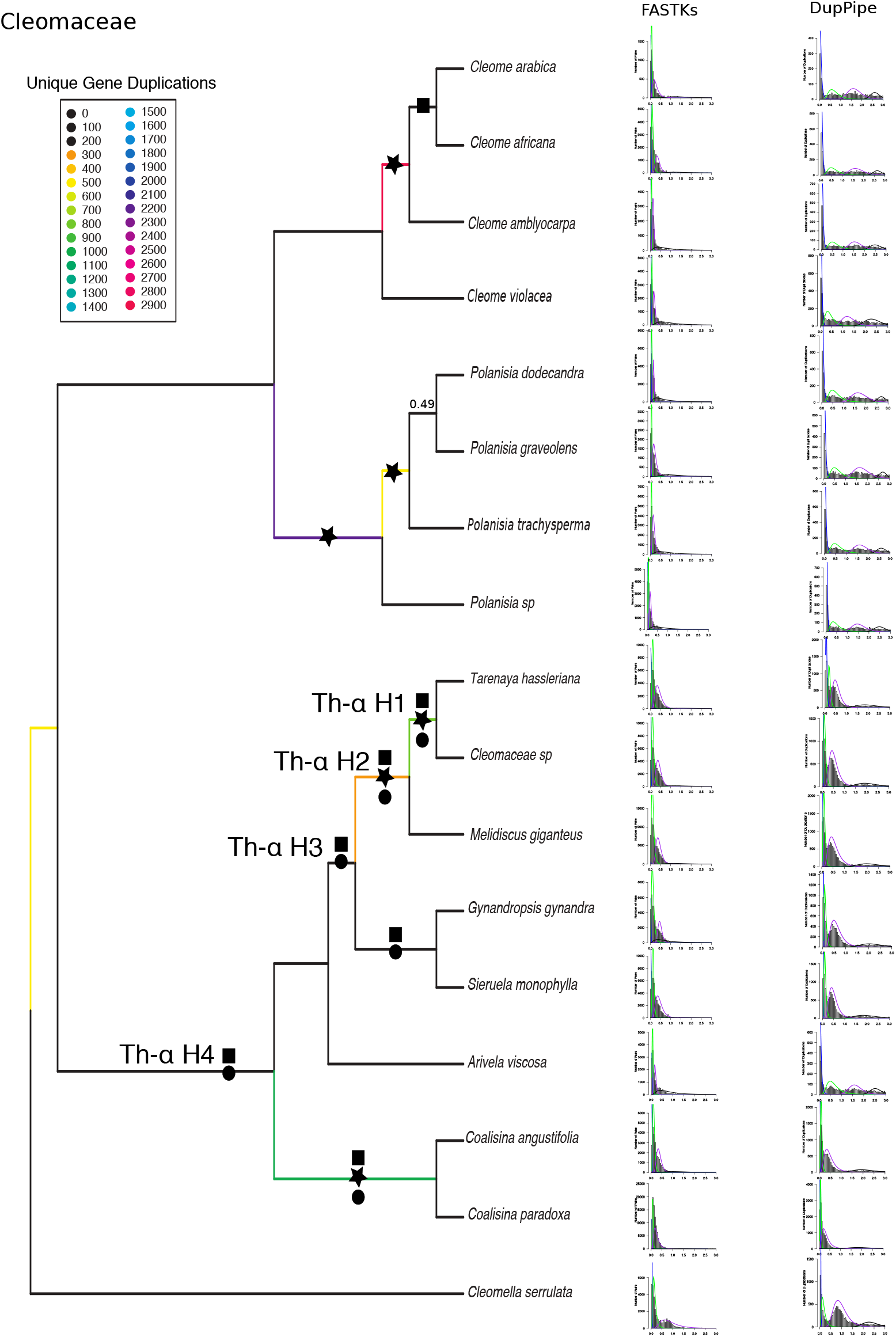
Coalescent-based species phylogeny and whole-genome duplication events of the Cleomaceae. Branches colored by number of unique gene duplications as determined by PUG (github.com/mrmckain/PUG). Black stars indicate WGD events identified by PUG, Black square indicate WGD events identified by FASTKs (McKain et al. 2016), and Black circles indicate WGD events identified by DupPipe (Barker et al. 2010). Ks plots using both FASTKs and DupPipe are placed next to their corresponding branch. Y-axes of Ks plots are not congruent, FASTKs measures number of pairs, while, DupPipe measures numbers of duplications. Support values are indicated if below 0.7 local posterior probability.

Both methods of Ks estimation provide support for the placement of Th-α with peaks ~ 0.4 for not only *T. hassleriana* and *Cleomaceae sp*, but also for *Melidiscus giganteus*, *Gynandropsis gynandra*, and *Sieruela monophylla*, suggesting that Th-α is shared across more than just *T. hassleriana* and *Cleomaceae sp* (Figure 3). However, we do not see evidence for this peak in *Arivela viscosa*, which is sister to the above species. As for the other three events, the story becomes more complicated. For many of them, when compared to *C. violacea* (which lacks evidence for Th-α from an analysis of its draft genome; Emery et al. 2018), one would conclude that there is no evidence for two of these novel events. Specifically, the one shared by *P. dodecandra*, *P. graveolens*, *P. trachysperma*, and *Polanisia sp.* or the one shared by *Cleome amblyocarpa*, *Cleome africana*, and *Cleome arabica*. However, the third potential event between *Coalisina paradoxa and Coalisina angustifolia* does have a signal for a WGD in the Ks plots (Figure 3).

Due to incongruence of results for the placement of Th-α, we divided potential placements into four hypotheses H1-H4 to test the age of ortholog divergence between taxa to the age of Th-α (Ks ~ 0.4). We find evidence that Th-α is shared with at least *T. hassleriana*, *Cleomaceae sp*, and *Melidiscus giganteus* and that Th-α occurred before the divergence between *Melidiscus giganteus* and *T. hassleriana* and around the same time as the divergence of *Gynandropsis gynandra* and *T. hassleriana* (Th-α H2; Figure 4A). We additionally compared the divergence between *A. viscosa* and *Gynandropsis gynandra* to the Ks values of the three species above along with *Sieruela monophylla*, *A. viscosa*, and *Gynandropsis gynandra*. We find that *A. viscosa* and *Gynandropsis gynandra* diverged more recently in time than Th-α and that, as in earlier Ks plots, *A. viscosa* lacks evidence for Th-α (Th-α H3; Figure 4B). This result is perplexing, and could indicate that the data from *A. viscosa* is either of poor quality or the genome itself has lost a large enough fraction of the duplicates that the signal for this event is not detected. To further test for the placement of Th-α we expanded our comparisons to include the ortholog divergence of *Coalisina angustifolia* and *T. hassleriana* as well as the divergence between *C. violacea* and *T. hassleriana* to test if the proposed independent WGD events between the two clades may in fact be a single event (Th-α H4; F**igure 4C**). We surprisingly recover both of these divergences to be of about the same age as Th-α, which would therefore have us conclude that Th-α is shared across this whole clade, and is not two separate events as illustrated in Figure 3. Comparison of ortholog divergence to Ks peaks for the two other identified WGD events using phylogenomics suggest that there is no other WGD event in the Cleomaceae (**Supp. Figure 6A & 6B**).

**Figure 4.**
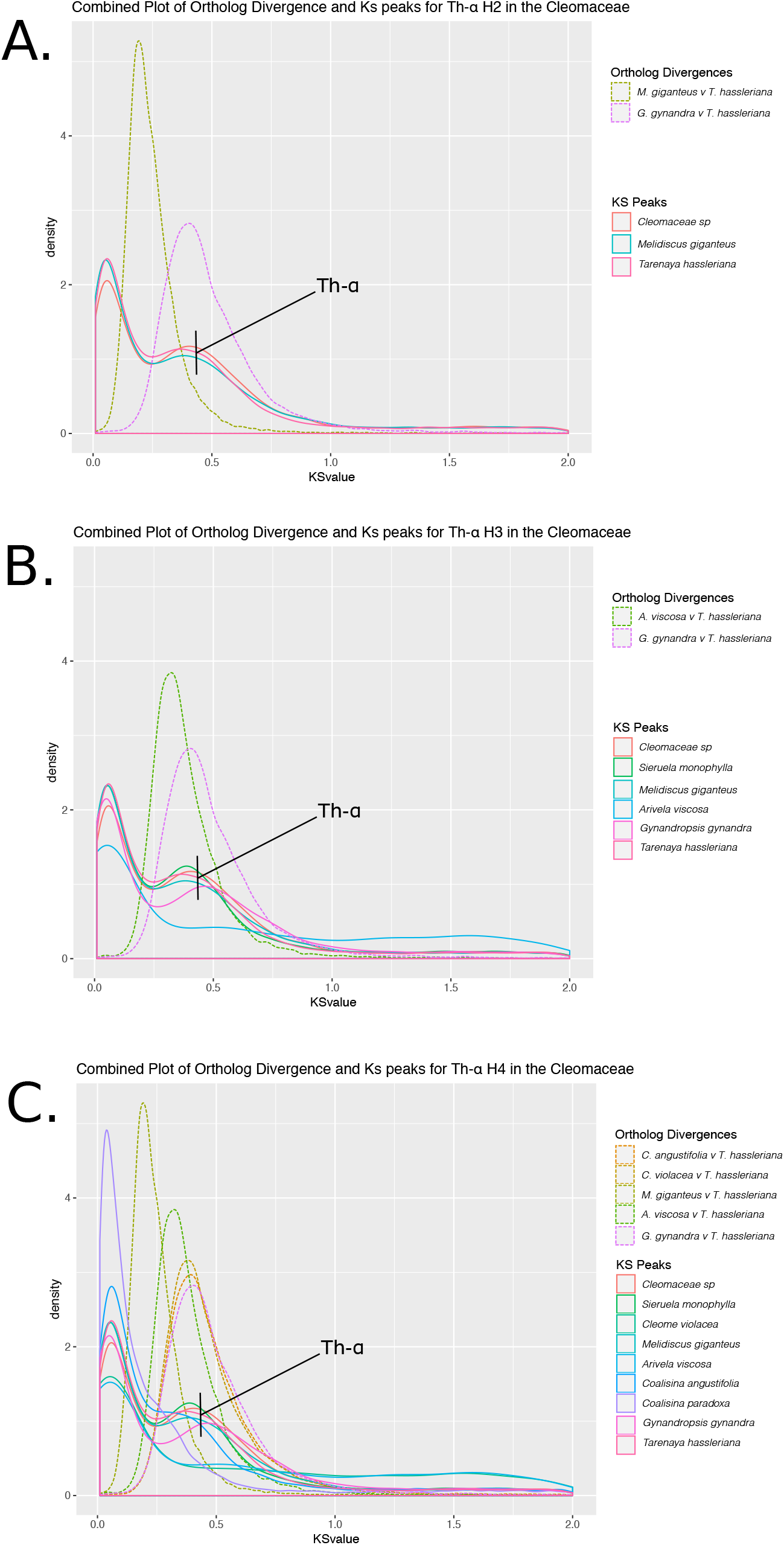
Comparison of ortholog divergences and Ks peaks of the Cleomaceae to test hypotheses of placement for Th-α. **(A)** Testing H2 by comparison of ortholog divergences of *Melidiscus giganteus* and *Gynandropsis gynandra* to *Tarenaya hassleriana* compared with Ks peaks of *Cleomaceae sp*, *Melidiscus giganteus*, and *Tarenaya hassleriana*. **(B)** Testing of H3 by comparison of ortholog divergence between *Arivela viscosa* and *Tarenaya hassleriana*, and *Gynandropsis gynandra* to *Tarenaya hassleriana* with Ks values of *Cleomaceae sp*, *Sieruela monophylla*, *Melidiscus giganteus, Arivela viscosa*, *Gynandropsis gynandra*, and *Tarenaya hassleriana*. **(C)** Testing the H4 hypothesis for placement of Th-α.

### Conflicting Evidence for WGD in the Capparaceae

In agreement with Lysak (2018), PUG recovers evidence for an independent WGD event in the Capparaceae, shared between a species of *Capparis* and another species of Capparaceae included in our analyses (Figure 5A). This event is also supported by Ks plots using FastKs, but not DupPipe, with a peak centered at Ks ~ 0.3 (Figure 5A). Ortholog divergences between members of the Capparaceae also show conflicting patterns. When comparing Ks values of *Boscia sp*, *Capparis fascicularis*, *Capparaceae sp*, and *Cadaba natalensis* from DupPipe to the ortholog divergence time between *Boscia sp* and *Capparis fascicularis* we find that the divergence between these two species occurred before the possible WGD event, agreeing with the PUG analysis. However, all four taxa share a peak in their Ks value, although their Ks plots from both analyses are not in agreement, providing conflicting results for the identification of this event. The divergences tested between *Boscia sp* and *Cadaba natalensis* as well as between *Capparis fascicularis* and *Cadaba natalensis* also occurred before the proposed event. However, the divergence between *Capparis fascicularis* and the misidentified species of Capparaceae, seems to have occurred at the same time as peak in Ks values (**Supp. Figure 7A**).

**Figure 5.**
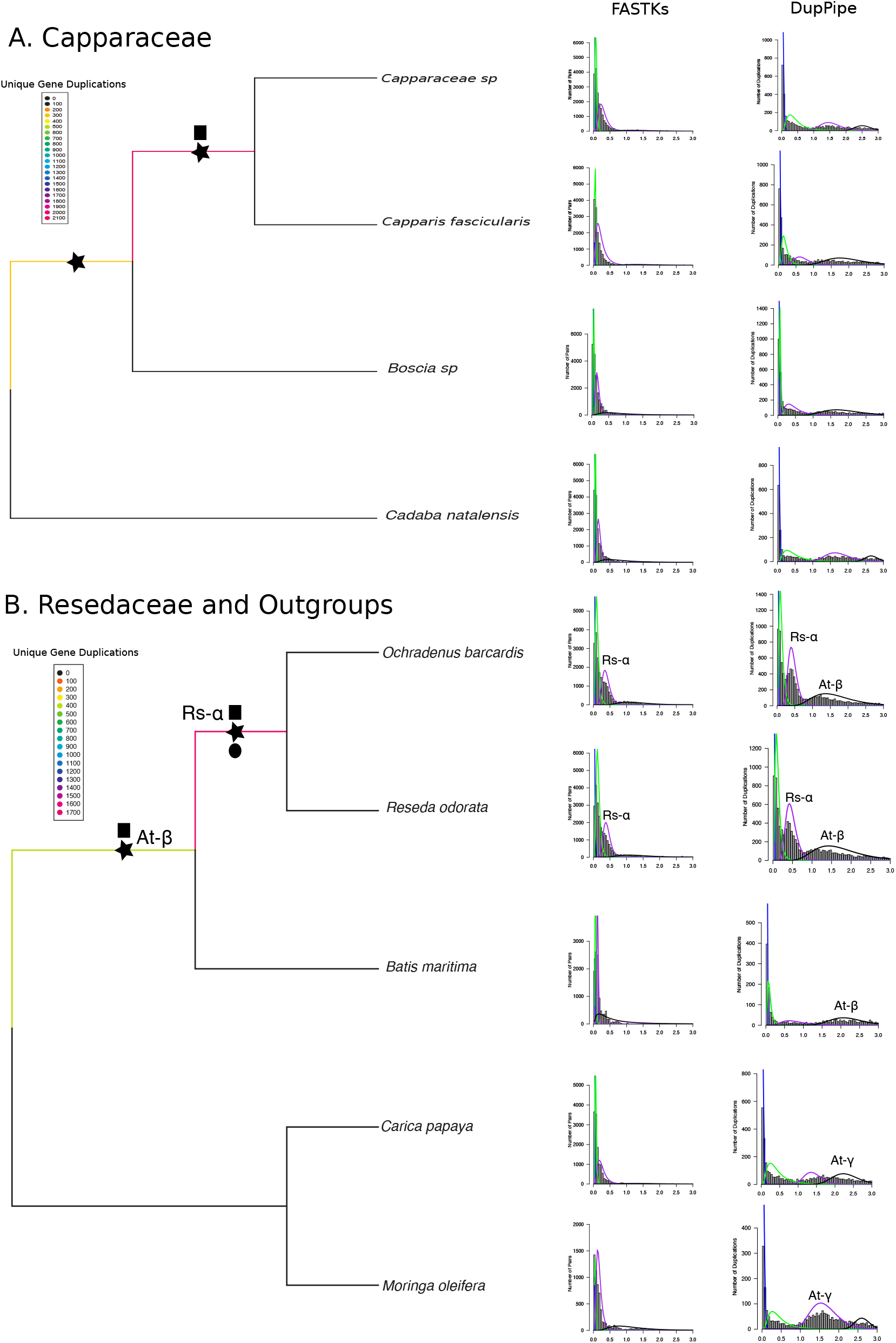
Coalescent-based species phylogenies and whole-genome duplication events of the **(A)** Capparaceae and **(B)** Resedaceae + Outgroups. Branches colored by number of unique gene duplications as determined by PUG (github.com/mrmckain/PUG). Black stars indicate WGD events identified by PUG, Black square indicate WGD events identified by FASTKs (McKain et al. 2016), and Black circles indicate WGD events identified by DupPipe (Barker et al. 2010). Ks plots using both FASTKs and DupPipe are placed next to their corresponding branch. Y-axes of Ks plots are not congruent, FASTKs measures number of pairs, while, DupPipe measures numbers of duplications. At-β and At-γ events noted above corresponding peaks in Ks plots. Support values are all above 0.7 local posterior probabilities.

### Novel WGD Event in the Resedaceae

When combining the Resedaceae (*Ochradenus barcardis* and *Reseda odorata*), Bataceae, Moringaceae, and Caricaceae families together, we excitedly find strong evidence for a Resedaceae specific WGD event in all three analyses with Ks plots indicating a peak ~ 0.4 (Figure 5B). Ortholog divergences seem to additionally support the proposal of this novel WGD between the samples of Resedaceae. Both samples (*Reseda odorata* and *Ochradenus barcardis*) share a Ks peak around ~ 0.4 which occurs before the divergence between these two samples and after the divergence between Resedaceae from *Batis maritima* (**Supp. Figure 7B**). In addition, we recover evidence for At-β using both PUG and DupPipe (Ks ~ 1.7; Figure 5B).

## DISCUSSION

Studies of the relationships within the Brassicales have either included many taxa but few genes (Hall et al. 2004; Hall 2008; Cardinal-McTeague et al. 2016), a few taxa and few genes (Rodman et al. 1998) or few taxa and many genes (Edger et al. 2015; Edger et al. 2018). In this study, we aim to find a balance of taxa and genes to present a well-supported chloroplast and nuclear phylogeny for the Brassicales, these both being in overall agreement with previous studies at the interfamilial and intrafamilial level (Edger et al. 2015; Cardinal-McTeague et al. 2016; Huang et al. 2016; Guo et al. 2017; Edger et al. 2018). Using the nuclear phylogeny, we highlight the difficulty in placing Th-α and identify possible novel events in the Cleomaceae, Capparaceae, and Resedaceae.

### Incongruences Between the Chloroplast and Nuclear Trees Across the Brassicaceae

Although relationships were congruent with previous analyses, we highlight the incongruence between the nuclear and chloroplast trees among the major lineages of the Brassicaceae, a well-documented pattern between these genomes (Beilstein et al. 2008; Huang et al. 2016; Nikolov et al. 2019; summarized in Figure 6). We find Lineage I sister to [Lineage III + Lineage II + Expanded Lineage II + Clade C] in the chloroplast tree and Lineage III sister to [Lineage I + Lineage II + Expanded Lineage II + Clade C]. Huang et al. (2016), using 113 low-copy nuclear genes from 55 Brassicaceae species, recovered a tree congruent with our nuclear tree with Lineage 1 sister to [Lineage III + Lineage II and Expanded Lineage II]. These relationships were also recovered by Nikolov et al. (2019), in their study using 79 species and 1,421 exons. Additionally, Guo et al. (2017) using 77 chloroplast genes from 53 samples, recovered a phylogeny in agreement with our chloroplast tree, with Lineage III sister to [Lineage I + Lineage II and Expanded Lineage II]. With additional taxon sampling, an increase in data, and using the same samples across analyses, we recover the congruent relationships, leaving us to conclude that the trees from these different genomes may never agree, potentially due to a complicated evolutionary history, such as ancient hybridization or introgression (Forsythe et al. 2018). These differences are important to consider when using the phylogeny to assess character evolution and divergence dating, as node ordering changes depending on which tree is used.

**Figure 6.**
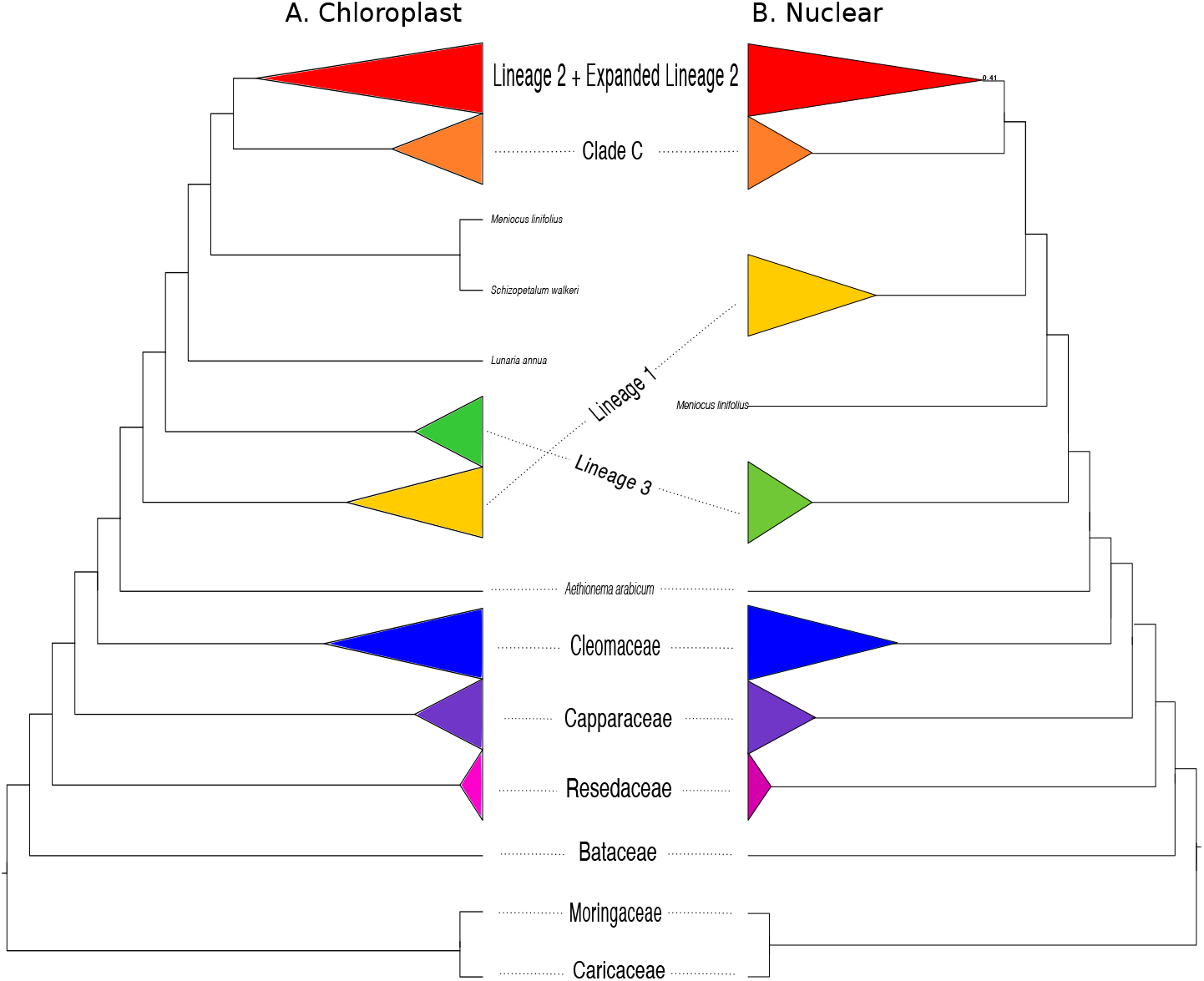
Comparison of **(A)** maximum likelihood whole-chloroplast phylogeny to **(B)** coalescent-based species phylogeny of the Brassicales. Major Lineages and clades of the Brassicaceae indicated. Support values are indicated if below 0.7 local posterior probabilities or 70 percent bootstrap support.

### Putative Placement of Th-α in the Cleomaceae

Previous studies have identified a WGD event unique to Cleomaceae (Th-α) using a variety of sources from syntenic regions to ESTs (Schranz & Mitchell-Olds 2006; Barker et al. 2009; reviewed in Bayat et al. 2018). However, the placement of Th-α within the Cleomaceae had yet to be confirmed. By using Ks plots to assess for signatures left behind in paralogs after WGD, phylogenetics using individual gene tree topologies, gene counts, and a known species tree, as well as ortholog divergences, we putatively place Th-α as shared between *Tarenaya hassleriana*, *Cleomaceae sp, Melidiscus giganteus, Gynandropsis gynandra*, and *Sieruela monophylla*, as well as *A. viscosa, Coalisina angustifolia*, and *Coalisina paradoxa* (Th-α H4; Figure 3). We have decided to include these last three species due to both the evidence from ortholog divergences and signatures in Ks plots that strongly suggest this event is shared with all species (Figures 3 & 4). However, there is a chance that two separate events occurred independently and that *A. viscosa* does indeed lack a WGD. Ks plots of all samples listed above, other than *A. viscosa*, identify a peak hovering over Ks ~ 0.4, agreeing with previous studies (Barker et al. 2009; van den Bergh et al. 2014) which first identified this peak in *Tarenaya hassleriana* followed by *Gynandropsis gynandra*. PUG, however, supports two separate events. One possibility for the difficulty in placing this event, may be due to the short branch lengths found within this clade (Figure 1) or that this event, like others in the order, is actually a triplication event and will be difficult to tease apart.

### Multiple WGD Events in the Cleomaceae?

In addition to identifying Th-α, we also report two possible additional events in the Cleomaceae, both of which are identified in the Brassicales and Cleomaceae specific analyses, but with much more support in the analysis of just Cleomaceae species. These WGD events are placed at common ancestors shared between: 1) *Cleome amblyocarpa, C. africana*, and *C. arabica* and 2) four species of *Polanisia* (Figure 3). Ks plots provide contrasting support for these events. Ks plots from FASTKs of *C. africana* and *C. arabica* show a small peak of duplicates hovering at Ks ~ 0.3, yet when the same data is run through DupPipe, there is no evidence of a WGD event. The Ks plots of *C. amblyocarpa* also provide conflicting evidence for this event. The Ks plot from FASTKs looks much more similar to that from *C. violacea*, which we know from sequencing its genome that it does not show evidence of any recent WGD event (Emery et al. 2018). While, the Ks plot from DupPipe of *C. amblyocarpa* complicates the story, with no clear peak identified. The second event, which is shared between four species of *Polanisia* is supported by a large number of unique gene duplications (2,200) using PUG, but is not supported by Ks plots from either FASTKs or DupPipe. The resulting plots look, again, more similar to *C. violacea*. Analyses of ortholog divergence between *C. amblyocarpa, C. africana*, and *C. arabica* also lack support for a WGD (Figure 6A) as do analyses between the four species of *Polanisia* (Figure 6B). To further test how WGD and C4 photosynthesis has evolved in this family, we suggest a study primarily focusing on Cleomaceae sampling. We know that C_4_ photosynthesis has evolved at least three times independently in Cleomaceae, specifically in (of the taxa sampled) *Gynandropsis gynandra* and *Coalisina angustifolia* (Bhide et al. 2014) with *Coalisina paradoxa* as a C3 – C4 intermediate in anatomy and physiology (Bhide et al. 2014). If our putative placement of Th-α is correct, then all of these samples share this event. Therefore, it will be interesting to investigate what the role of polyploidy, and more specifically Th-α, is in character evolution in this group.

### Novel WGD Events in the Capparaceae and Resedaceae

Although just two samples are included in our analysis, we recover some support for an event between at least one species of *Capparis* and a misidentified species of Capparaceae (Figure 5A). Due to this possible identification error, inconclusiveness from Ks plots, and no support in comparison between ortholog divergence and Ks peaks, this event, although supported by many unique duplicates in the PUG analysis, should be interpreted carefully. However, Lysak (2018), using chromosome counts, also proposed that Capparaceae had a unique event, making this an intriguing event to further investigate. It should be noted, however, that chromosome counts alone may be misleading in concluding that a WGD event has occurred (Evans et al. 2017). Alternatively, looking at just Ks plots for this group, it is still difficult to ascertain if there is a unique event. Capers are typically much more woody than the others plants we sampled and therefore have longer generation times, which needs to be accounted for when interpreting peaks derived from Ks plots. There is also a lack of agreement between Ks plots derived using FASTKs and DupPipe which estimate Ks values in different ways (pairwise Ks estimates in FASTKs versus estimates of Ks at nodes in gene trees in DupPipe), further confounding evidence for either a presence or absence of a Capparaceae-specific event. Between information presented by Lysak (2018) and the evidence presented here, this possible event certainly warrants additional study.

A separate WGD event in the Resedaceae was also hypothesized by Lysak (2018), here we find good evidence to support its presence. This is one of the few events recovered with consensus between Ks plots (from both FASTKs and DupPipe), phylogenetics, and ortholog divergences (Figure 4B & **Supp. Fig. 7B**). Therefore, we are confident in naming this event as Rs-α. The sister families, Caricaceae and Moringaceae, show no evidence of unique WGD events, which is in agreement with the recent whole-genome sequencing of *Moringa oleifera* (Chang et al. 2018). When Tian et al. (2015) compared the papaya genome, which shows no evidence of a (recent) WGD (Ming et al. 2008), to their newly sequenced genome of *Moringa oleifera*, they too concluded that Moringaceae did not experience a family-specific genome duplication. Although we only surveyed two Resedaceae species, we feel this event is well supported and warrants additional sampling and investigation.

### Methodological Challenges with Placing WGD Events; Sampling Matters

Currently three types of methods are used to detect WGD; Ks plots to assess for signatures left behind in paralogs after WGD, identification of retained duplicate blocks in a genome, and phylogenetics using individual gene tree topologies, gene counts, and a known species tree, with Ks plots and phylogenomics being the most approachable. All three of these methods however, have their limitations in identifying WGD events. As others have noted, and we have done here too, using a combination of approaches to test hypotheses helps to reduce the chance of proposing events that may not exist and simultaneously provides multiple lines of evidence for those events that are recovered.

Recently, there has been an abundance of papers highlighting the difficulties and complexities of determining WGD events across the tree of life (Conover et al. 2018; Tiley et al. 2018; Li & Barker 2019; Li et al. 2019; Nakatani & McLysaght 2019; Zwaenepoel & Van de Peer 2019; Zwaenepoel et al. 2019). We add another dimension to this conversation by demonstrating that the different taxonomic levels in which we sampled, such as the order or family made a difference in support of known events (i.e., the Brassiceae triplication). Recent research has demonstrated that differences in taxonomic sampling and taxon occupancy in data matrices can influence the inference of WGDs, particularly if adding taxa decreases taxon occupancy in gene families (Yang et al. 2015; Li et al. 2018; Li & Barker 2019; Zwanepoel & Van de Peer 2019). Testing for WGD events across the Brassicales phylogeny also led to less certain topologies, therefore when filtering for nodes with only high bootstrap support to count duplicates, signals of WGD may be missed. To account for this, and increase taxon and gene family occupancy in our datasets, we reduced sampling to just the family level. However, at each level of analysis, we had to choose an arbitrary cut-off for the number of duplicates that we felt were sufficient to infer a WGD event, a documented criticism of these types of methods (Zwaenepoel & Van de Peer, 2019). Many authors also note that it is important to consider heterogeneity in substitution rates (Barker et al. 2009; Yang et al. 2015) as well as variation in the duplication and loss rate across the species tree when testing for WGD events (Li et al. 2018; Zwaenepoel & Van de Peer 2019).

Although our Ks based inferences of WGDs were largely consistent with the phylogenomic inferences, there were some differences among the approaches. FASTKs and DupPipe use different estimates of Ks that likely produced the observable differences in their respective Ks plots. FASTKs uses a pairwise approach to estimate Ks values (github.com/mrmckain/FASTKs; McKain et al. 2016), whereas DuPipe estimates Ks values from nodes of gene trees (Barker et al. 2010). The difference in Ks estimates from these types of approaches was previously explored by Tiley et al. (2018), and the observed differences in peaks of duplications between the two different methods is consistent with simulations (Tiley et al. 2018). The node-based estimates of Ks from DupPipe often yielded apparently sharper peaks in putative WGDs with overall lower numbers of duplications because of the difference in number of nodes vs pairwise comparisons. However, the results of both approaches were largely consistent after close inspection. Perhaps more confounding for Ks analyses is the interpretation of mixture models to identify putative peaks associated with a WGD. Mixture models, which are typically fit to the distribution of duplicates, tend to overestimate the number of true peaks (Naik et al. 2007; Tiley et al. 2018; Zwaenepoel et al. 2019). Using the two different methods as we did here, and across multiple species, allowed us to evaluate and compare putative peaks from different analyses to identify the expected signatures of WGDs. Further, paralogs from WGDs tend to be expressed more than those resulting from tandem duplications (Casneuf et al. 2006), using transcriptome data, as we did here, may actually yield data that is more enriched for WGD duplicates than using (fragmented) genomic data. Therefore using transcriptome data, as shown by Tiley et al. (2018), may actually improve our success in detecting WGD events.

Overall, the Brassicales are an excellent group of plants to compare methods of WGD identification because of the wealth of genomic data available and known events. With many chromosome level genomes available, analyses based on syntenty, which seem to be regarded as most reliable in detecting these events (Nakatani & McLysaght 2019), can be used as controls for comparing WGD methods. Sequenced genomes, which are placed throughout the Brassicales, provide strong evidence for taxa in which we know do not have recent WGD events (i.e., *Cleome violacea* and *Carica papaya*) and taxa that do show evidence for recent WGD events (i.e. *Arabidopsis thaliana* and many *Brassica* crops). Because of these resources, we have calibration points that allow us to verify results when testing for novel events. Perhaps this group of plants, combined with recent insights on difficulties in placing WGD events, will help in furthering the development of innovative methods in describing and identifying WGDs.

## METHODS

### Taxon Sampling

Sampling of 74 species of 57 genera across the Brassicales spanned seven families (Brassicaceae, Cleomaceae, Capparaceae, Resedaceae, Bataceae, Moringaceae, and Caricaceae), with a focus on the Brassicaceae (48 taxa) and Cleomaceae (17 taxa) (**Supp. Table 2)**. Seeds were grown at the University of Missouri - Columbia or the University of Alberta in a sterile growth chamber environment. At maturity, but before flowering, leaf tissue was collected for both RNA and DNA extraction.

### DNA and RNA Isolation and Sequencing

DNA was extracted from leaf tissue for 69 of the 74 taxa using the Qiagen DNeasy Plant kit (Qiagen, Germantown, MD, USA). To increase yield, slight modifications to the manufacturer’s protocol included increasing lysis buffer incubation time to one hour and using 25 µl of buffer to elute the final sample. TruSeq library preparation (Illumina) and sequencing on a NextSeq (Illumina) took place at the University of Missouri-Columbia resulting in 2 × 150 bp reads. At the University of Missouri, RNA sampling of leaf tissue was collected and immediately flash frozen using liquid nitrogen. For 38 samples, RNA was isolated using the ThermoFisher Invitrogen PureLink RNA mini kit (Invitrogen, Carlsbad, CA, USA) followed by TruSeq library preparation (Illumina) and sequencing on the NextSeq (Illumina) resulting in 2 × 75 bp reads (**Supp. Table 3**). For 16 samples, RNA was again isolated using the ThermoFisher Invitrogen PureLink RNA mini kit (Invitrogen, Carlsbad, CA, USA), however sequencing took place on an HiSeq instrument resulting in 2 × 100 bp reads (**Supp. Table 3**). For 17 samples, RNA was sequenced on an HiSeq instrument for 2 × 100 bp reads, but used the Qiagen RNeasy Plant Kit (Qiagen, Germantown, MD, USA) for RNA isolation (**Supp. Table 3**). Lastly, two samples were isolated again using the ThermoFisher Invitrogen PureLink RNA mini kit (Invitrogen, Carlsbad, CA, USA), but were sequenced on a HiSeq for 2 × 250 bp reads (**Supp. Table 3**). All sequencing and library preparation for the above samples was performed by the University of Missouri DNA Core Facility. At the University of Alberta, one sample, *Cleomella serrulata* tissue was pooled from leaves, apical meristematic tissue, and floral tissue of different developmental stages including small, medium, and large buds, and open flowers from two plants. All the collected tissue was flash frozen in liquid Nitrogen and kept at −80 °C to avoid RNA degradation. Total RNA was extracted with RNeasy plant MiniKit (Qiagen, Hilden, Germany) following the manufacturer’s protocol, then treated with DNAse I (New England Biolabs, Ipswich, USA) for 30 min at 37 °C to remove residual DNA from the total RNA. Sequencing was then conducted by Plate-forme d’Analyses Génomique de l’ Université Laval by purifying mRNA from 3 µg of total RNA, then fragmenting and converting it to double-stranded cDNA using the Illumina TruSeq RNASeq library preparation kit following Illumina’s guidelines.

### Chloroplast Assembly, Alignment, and Phylogenomics

For analysis of just two chloroplast genes, matK and ndhF, we included 91 taxa from the study by Hall (2008). The two chloroplast genes were annotated and extracted from chloroplast sequences using Geneious v8.1.9 (Kearse et al. 2012). For one taxon, *Batis maritima*, we were unable to annotate and extract ndhF. Alignment of resulting genes was performed in MAFFT v7 (Katoh 2002) and cleaned using phyutility v2.7.1 (Smith and Dunn 2008) with the parameter *-clean 0.5*. For maximum likelihood (ML) phylogenetic inference, RAxML v8 (Stamatakis 2014) was run with a separate partition for each gene, GTRGAMMA as the model, and 1000 bootstrap replicates.

To assemble whole chloroplasts, Fast-Plast v1.2.8 was used (McKain and Wilson 2017). This method utilizes Trimmomatic v0.35 (Bolger et al. 2014) to clean the reads of adaptors using a Phred score of 33, Bowtie2 v2.3.4.3 (Langmead et al. 2012) to separate chloroplast reads by mapping them to a reference database of Angiosperm chloroplasts, followed by both SPAdes v3.13.0 (Bankevich et al. 2012) and afin to assemble reads. For 13 samples which would not assemble with the default options, the *--subsample* option yielded successful assemblies (**Supp. Table 2**). For three samples, *Polanisia dodecandra*, *Farsetia aegyptia*, and *Cardamine hirsuta*, we were only able to obtain partial regions of the chloroplast genome, and therefore they were not used in downstream analyses (**Supp. Table 2**). Following assembly, MAFFT v7 (Katoh 2002) was used to align the LSC region, the SSC region, and one copy of the IR. Alignments were cleaned using phyutility v2.7.1 (Smith and Dunn 2008) with the parameter *-clean 0.5*. Finally for ML phylogenomic inference, RAxML v8 (Stamatakis 2014) was run with partitions for each region, GTRGAMMA as the model, and 1000 bootstrap replicates.

### Transcriptome Assembly, Alignment, and Phylogenomics

For transcriptome analyses, reads were trimmed with trimmomatic v0.35 (Bolger et al. 2014) using the parameters *SLIDINGWINDOW:4:5*, *LEADING:5*, *TRAILING:5*, and *MINLEN:25* followed by assembly using Trinity v2.2 (Grabherr et al. 2011). The resulting transcriptomes were checked for completeness using BUSCO v3 (Simão et al. 2015; Waterhouse et al. 2017) and compared to the Embryophyta database. Transcriptomes were translated to protein sequences and coding regions were predicted using TransDecoder v3.0 (github.com/TransDecoder/TransDecoder). Finally, orthology was inferred using OrthoFinder v.2.2.6 (Emms & Kelly 2018), first with the parameters *-S diamond* (Buchfink et al. 2015), then for a second time with the parameters *-M msa -ot* for multiple sequence alignments and only trees. Using custom scripts, alignments were filtered for 80% taxon occupancy (github.com/MU-IRCF/filter_by_ortho_group) and alignment quality, allowing for only 40% gaps (github.com/MU-IRCF/filter_by_gap_fraction). To estimate gene trees using ML inference, RAxML v8 was used (Stamatakis 2014) followed by PhyloTreePruner v1.0 (Kocot et al. 2013) to remove any potentially remaining paralogous genes. Since alignments had previously been filtered for taxon occupancy, a cutoff of 10 was used for the minimum number of taxa required to keep a group. Alignments passing this threshold were then used to estimate final gene trees using RAxML v8 (Stamatakis 2014). Species tree estimation was then performed using ASTRAL-III v.5.6.1 (Zhang et al. 2018). Analyses were performed on all samples (Brassicales) and at the family level (Brassicaceae, Cleomaceae, Capparaceae, [Resedaceae + Bataceae + Moringaceae + Cariacacae]; **Supp. Table 1**).

### Whole-Genome Duplication

To estimate the phylogenetic placement of whole-genome duplications, PUG v2.1 (github.com/mrmckain/PUG) was used to query putative paralogs over multiple gene trees using the estimated ASTRAL-III tree as the input species tree. For each analysis, we used the original ML gene trees before running them through PhyloTreePruner (i.e., gene trees with all duplicates retained), the ASTRAL-III tree (rooted and with bootstraps removed), and parameters *-- estimate_paralogs* and *--outgroups Carica_papaya,Moringa_oleifera* as input. Output duplicate gene counts were used only for those nodes with bootstrap values of 80 percent or better.

As another confirmation of duplication events, we constructed histograms giving the distribution of the synonymous rate of divergence (Ks) between paralogs in each transcriptome. This method allows for the potential identification of peaks in the distribution that may be indicative of a WGD event. The position of the peak along the Ks axis provides an estimate of time when the event occurred. Typically the peak closest to time zero (or Ks ~ 0) corresponds to recent tandem duplicates, not relevant to WGD events. Plots of Ks distributions were made for all taxa using FASTKs v1.1 (github.com/mrmckain/FASTKs) as described in McKain et al. (2016) and DupPipe following Barker et al. (2010). Following Ks analyses, R v3.5.1(R Core Team 2018), was used to estimate normal mixture models for Ks values using mclust v.5.0.2 (Fraley and Raftery 2002; Fraley et al. 2012). To test for the best number of peaks to explain the data, we tested one to four components for each mixture model. We then picked the one with the lowest Bayesian information criterion (BIC) score as the best fit (**Supp. Table 4**). Although for most taxa four components had the lowest score, we emphasize that this does not mean that there are four WGD duplication events.

To further test for phylogenetic placement of WGD events, ortholog divergence was estimated using OrthoPipe as described in Barker et al. (2010). Using the estimated ortholog divergence and DupPipe Ks estimates, we are able to bookend the position of potential events by comparing when species diverged to the age of an estimated WGD event. If the ortholog divergence between pairs of species is older (larger Ks value) than the estimated age of a WGD event, one can conclude that those species do not share the event, however, if ortholog divergence between species is younger than the WGD, species do share the proposed event.

## Accession Numbers

Sequence data from this article can be found in the NCBI SRA data libraries under BioProject accession number PRJNA542714. Individual BioSample accession numbers can be found in **Supplemental Table 1**.

## Supplemental Data

The following materials are available in the online version of this article.

**Supplemental Figure 1.** Current phylogenetic relationships between the 17 families of the Brassicales.

**Supplemental Figure 2.** BUSCO analysis of de novo transcriptomes.

**Supplemental Table 1.** Taxon sampling, accessions, and additional analysis information.

**Supplemental Figure 3.** Maximum Likelihood Phylogeny of the Brassicales using two chloroplast genes, MatK and NdhF.

**Supplemental Figure 4.** Maximum Likelihood Whole-Chloroplast Phylogeny of the Brassicales.

**Supplemental Figure 5.** Brassicaceae Ks plots using both FASTKs (McKain et al. 2016) and DupPipe (Barker et al. 2010).

**Supplemental Figure 6.** Additional Ortholog divergences and Ks peaks of the Cleomaceae.

**Supplemental Figure 7.** Ortholog divergences and Ks peaks of the (A) Capparaceae and (B) Resedaceae + Outgroups.

**Supplemental Table 2.** Orthogroups retained for each analysis.

**Supplemental Table 3.** RNA and DNA extraction method, library preparation method, sequencing method, read size, and raw read numbers.

**Supplemental Table 4.** BIC scores for 1-4 components for both FASTKs (McKain et al. 2016) and DupPipe (Barker et al. 2010).

## ACKNOWLEDGEMENTS

We would like to our funding sources including NSF grants IOS-1339156 and EF-1550838 to MSB, NSF IOS-1811784 to PDB, NIH PERT K12GM000708 to BS, and NSF DEB-1146603 to JCP. We also thank the University of Missouri DNA Core Staff - Nathan Bivens, Ming-Yi Zhou, and Karen Bromert for their patience and assistance in getting successful libraries for samples which we had very little DNA or RNA for and the University of Missouri Research Computing Support Services (RCSS); specifically Jake Gotburg, for his assistance in running large jobs across the Lewis cluster. We also thank Barb Sonderman for her assistance in helping to keep many of these plants alive. Finally, we thank Sarah Unruh and our anonymous reviewers for providing feedback in improving this manuscript.

## AUTHOR CONTRIBUTIONS

MEM, JCP, GCC, JCH, PPE, and MES designed the project. MEM and JMB analyzed the data. PDB, CAB, AH, MSB, BS, and MRM assisted with processing the data. MEM, JMB, JDW, and WTD prepared RNA and DNA for sequencing. MEM, JMB, JDW, WTD, and IA planted, sampled, and phenotyped the plant materials. MEM wrote the manuscript.

---

Supp. Figure 1. Current understanding of the relationships between the 17 families of the Brassicales (APG IV). * indicates branch support between 50-80%, all other branches have greater than 80% support.

Supp. Figure 2. BUSCO analysis of de novo transcriptomes. Legend indicates the percent of genes that are complete and single copy (light blue), complete and duplicate (dark blue), fragmented (yellow), and missing (red) in de novo transcriptomes.

Supp. Figure 3. Maximum likelihood phylogeny of the Brassicales using two chloroplast genes, MatK and NdhF. Support values are indicated if below 70 percent bootstrap support. * next to taxa indicate those whose placement are not sister with samples from Hall (2008).

Supp. Figure 4. Maximum likelihood whole-chloroplast phylogeny of the Brassicales. Support values are indicated if below 70 percent bootstrap support.

Supp. Figure 5. Ks plots of the Brassicaceae using both FASTKs (McKain et al. 2016) and DupPipe (Barker et al. 2010). Whole-genome duplication events, At-α and the Brassiceae triplication (T) event are noted above corresponding peaks.

Supp. Figure 6. Ortholog divergences and Ks peaks of the Cleomaceae. **(A)** Ortholog divergences between *C. amblyocarpa* and *C. africana, C. arabica*, and *C. violacea* and between *C. violacea* and *C. africana* to test placement of potential novel WGD event. **(B)** Ortholog divergences between *Polanisia sp.* and *C. violacea, Polanisia sp.* and *P. dodecandria, P. trachysperma* and *Polanisia sp.*, and between *P. trachysperma* and *P. dodecandria* to test for placement of the second potential novel WGD event.

Supp. Figure 7. Ortholog divergences and Ks peaks of the **(A)** Capparaceae and **(B)** Resedaceae + Outgroups. Proposed Resedaceae whole-genome duplication event indicated.

Supp. Table 1. Taxon sampling, seed accessions and the collections they are from, additional analysis information, and SRA numbers for both RNA and genome survey sequencing (GSS) raw reads. SSC = small single copy, LSC= large single copy, IR = inverted repeat.

Supp. Table 2. Orthogroups retained for each analysis.

Supp. Table 3. RNA and DNA extraction method, library preparation method, sequencing method, read size, and raw read numbers.

Supp. Table 4. BIC scores for 1-4 components for both FASTKs (McKain et al. 2016) and DupPipe (Barker et al. 2010) Ks plots.

## REFERENCES

APG IV. (2016). An update of the Angiosperm Phylogeny Group classification for the orders and families of flowering plants: APG IV. Botanical Journal of the Linnean Society 181(1): 1–20.

Arabidopsis Genome Initiative. (2000). Analysis of the genome sequence of the flowering plant *Arabidopsis thaliana*. Nature, 408(6814), 796.

Barker M.S., Vogel, H., and Schranz, M. E. (2009). Paleopolyploidy in the Brassicales: analyses of the Cleome transcriptome elucidate the history of genome duplications in Arabidopsis and other Brassicales. Genome Biology and Evolution, 1: 391–399.

Barker, M. S., Dlugosch, K. M., Dinh, L., Challa, R. S., Kane, N. C., King, M. G., and Rieseberg L. H. (2010). EvoPipes. net: bioinformatic tools for ecological and evolutionary genomics. Evolutionary Bioinformatics, 6: EBO–S5861.

Bankevich, A., Nurk, S., Antipov, D., Gurevich, A.A., Dvorkin, M., Kulikov, A.S., Lesin, V.M., Nikolenko, S.I., Pham, S., Prjibelski, A.D., Pyshkin, A.V., Sirotkin, A.V., Vyahhi, N., Tesler, G., Alekseyev, M.A., and Pevzner, P.A. (2012). SPAdes: a new genome assembly algorithm and its applications to single-cell sequencing. Journal of Computational Biology, 19(5): 455–477.

Bayat, S., Schranz, M. E., Roalson, E. H., and Hall, J.C. (2018). Lessons from Cleomaceae, the sister of crucifers. Trends in Plant Science 23: 808–821

Beilstein, M.A., Al-Shehbaz, I.A., and Kellogg, E.A. (2006). Brassicaceae phylogeny and trichome evolution. American journal of botany 93(4): 607–619.

Beilstein, M.A., Al-Shehbaz, I.A., Mathews, S., and Kellogg, E.A. (2008). Brassicaceae phylogeny inferred from phytochrome A and ndhF sequence data: tribes and trichomes revisited. American Journal of Botany, 95(10), pp.1307–1327.

Bhide A., Schliesky, S., Reich, M., Weber, A.P., and Becker, A. (2014). Analysis of the floral transcriptome of *Tarenaya hassleriana* (Cleomaceae), a member of the sister group to the Brassicaceae: towards understanding the base of morphological diversity in Brassicales. BMC Genomics 15(1): 140.

Bolger, A. M., Lohse, M., and Usadel, B. (2014). Trimmomatic: a flexible trimmer for Illumina sequence data. Bioinformatics 30(15): 2114–2120.

BrassiBase. Tools and biological resources for Brassicaceae character and trait studies. https://brassibase.cos.uni-heidelberg.de. Accessed 24 Mar 2019.

Brock, K.C. (2014). Tracking the Evolutionary History of Development Genes: Implications for the Diversification of Fruits and Flowers in the Brassicaceae and Cleomaceae. Master’s thesis. University of Alberta, Edmonton, Alberta, Canada.

Buchfink, B., Xie, C., and Huson, D.H. (2015). Fast and sensitive protein alignment using DIAMOND. Nature Methods 12(1): 59.

Cardinal-McTeague W., Sytsma, K.J., and Hall, J.C. (2016). Biogeography and diversification of Brassicales: A 103 million year tale. Mol Phylogenetics and Evolution 99: 204–224

Casneuf, T., De Bodt, S., Raes, J., Maere, S., and Van de Peer, Y. (2006). Nonrandom divergence of gene expression following gene and genome duplications in the flowering plant *Arabidopsis thaliana*. Genome Biology 7(2): R13.

Chang, Y., Liu, H., Liu, M., Liao, X., Sahu, S. K., Fu, Y., Song, B., Cheng S., Kariba, R., Muthemba, S., Hendre, P.S., Mayes, S., Ho, W.K., Yssel, A.E.J., Kendabie, P., Wang, S., Li, L., Muchugi, A., Jamnadass, R., Lu, H., Peng, S., Van Deynze, A., Simons, A., Yana-Shapiro, H., Van de Peer, Y., Xu, X., Yang, H., Wang, J., Liu, X. (2018). The draft genomes of five agriculturally important African orphan crops. GigaScience 8(3): giy152.

Cheng, S., van den Bergh, E., Zeng, P., Zhong, X., Xu, J., Liu, X., Hofberger, J., de Bruijn, S., Bhide, A.S., Kuelahoglu, C., Bian, C., Chen, J., Fan, G., Kaufmann, K., Hall, J.C., Becker, A., Bräutigam, A., Weber, A.P.M., Shi, C., Zheng, Z., Li, W., Lv, M., Tao, Y., Wang, J., Zou, H., Quan, Z., Hibberd, J.M., Zhang, G., Zhu, X.G., Xu, X., and Schranz, M.E. (2013). The *Tarenaya hassleriana* genome provides insight into reproductive trait and genome evolution of crucifers. The Plant Cell, 25(8), 2813–2830.

Conover, J. L., Karimi, N., Stenz, N., Ané, C., Grover, C. E., Skema, C., Tate, J.A., Wolff, K., Logan, S.A., Wendel, J.F., and Baum, D. (2018). A Malvaceae mystery: A mallow maelstrom of genome multiplications and maybe misleading methods?. Journal of integrative plant biology 61(1): 12–31.

Duarte, J. M., Wall, P. K., Edger, P. P., Landherr, L. L., Ma, H., Pires, J. C., and Leebens-Mack, J. (2010). Identification of shared single copy nuclear genes in *Arabidopsis*, *Populus*, *Vitis* and *Oryza* and their phylogenetic utility across various taxonomic levels. BMC Evolutionary Biology, 10(1): 61.

Dunn, C.W., Hejnol, A., Matus, D.Q., Pang, K., Browne, W.E., Smith, S.A., Seaver, E., Rouse, G.W., Obst, M., Edgecombe, G.D., and Sørensen, M.V. (2008). Broad phylogenomic sampling improves resolution of the animal tree of life. Nature, 452(7188), p.745.

Edger P. P., Heidel-Fischer, H.M., Bekaert, M., Rota, J., Glöckner, G., Platts, A.E., Heckel, D.G., Der, J. P., Wafula, E.K., Tang, M., Hofberger, J.A., Smithson, A., Hall, J.C., Blanchette, M., Bureau, T.E., Wright, S.I., dePamphilis, C.W., Schranz, M.E., Barker, M.S., Conant, G.C., Wahlberg, N., Vogel, H., Pires, J.C., and Wheat, C.W. (2015). The butterfly plant arms-race escalated by gene and genome duplications. Proceedings of the National Academy of Sciences 112(27): 8362–8366.

Edger, P.P., Hall, J.C., Harkess, A., Tang, M., Coombs, J., Mohammadin, S., Schranz, M.E., Xiong, Z., Leebens-Mack, J., Meyers, B.C., Sytsma, K.J., Koch, M.A., Al-Shehbaz, I.A., and Pires J.C. (2018). Brassicales phylogeny inferred from 72 plastid genes: A reanalysis of the phylogenetic localization of two paleopolyploid events and origin of novel chemical defenses. American journal of botany, 105(3), pp.463–469.

Emery, M., Willis, M.M.S., Hao, Y., Barry, K., Oakgrove, K., Peng, Y., Schmutz, J., Lyons, E., Pires, J.C., Edger, P.P., and Conant, G.C. (2018). Preferential retention of genes from one parental genome after polyploidy illustrates the nature and scope of the genomic conflicts induced by hybridization. PLoS genetics, 14(3), p.e1007267.

Emms, D. M., and Kelly, S. (2015). OrthoFinder: solving fundamental biases in Whole-genome comparisons dramatically improves orthogroup inference accuracy. Genome Biology 16(1): 157.

Emms, D.M. and Kelly, S. (2018). OrthoFinder2: fast and accurate phylogenomic orthology analysis from gene sequences. bioRxiv.

Endress, P. K. (1992). Evolution and floral diversity: the phylogenetic surroundings of *Arabidopsis* and *Antirrhinum*. International Journal of Plant Sciences 153(3, Part 2): S106–S122.

Evans, B. J., Upham, N. S., Golding, G. B., Ojeda, R. A., and Ojeda, A. A. (2017). Evolution of the largest mammalian genome. Genome biology and evolution, 9(6), 1711–1724.

Forsythe, E.S., Nelson, A.D., and Beilstein, M.A. (2018). Biased gene retention in the face of massive nuclear introgression obscures species relationships. bioRxiv, p.197087.

Fraley, C., and Raftery, A.E. (2002). Model-based clustering, discriminant analysis, and density estimation. Journal of the American statistical Association 97(458): 611–631.

Fraley, C., Raftery, A.E., and Scrucca, L. (2012). Normal mixture modeling for model-based clustering, classification, and density estimation. Department of Statistics University of Washington, 23, 2012.

Grabherr, M.G., Haas, B.J., Yassour, M., Levin, J.Z., Thompson, D.A., Amit, I., Adiconis, X., Fan, L., Raychowdhury, R., Zeng, Q., Chen, Z., Mauceli, E., Hacohen, N., Gnirke, A., Rhind, N., di Palma, F., Birren, B.W., Nusbaum, C., Lindblad-Toh, K., Friedman N., and Regev, A. (2011). Full-length transcriptome assembly from RNA-Seq data without a reference genome. Nature biotechnology, 29(7), p.644.

Guo, X., Liu, J., Hao, G., Zhang, L., Mao, K., Wang, X., Zhang, D., Ma, T., Hu, Q., Al-Shehbaz, I.A. and Koch, M.A. (2017). Plastome phylogeny and early diversification of Brassicaceae. BMC genomics, 18(1), p.176.

Hall, J. C., Sytsma, K. J., and Iltis, H.H. (2002). Phylogeny of Capparaceae and Brassicaceae based on chloroplast sequence data. American Journal of Botany 89(11): 1826–1842.

Hall J. C., Iltis, H. H., Sytsma, K. J. (2004). Molecular phylogenetics of core Brassicales, placement of orphan genera Emblingia, Forchhammeria, Tirania, and character evolution. Systematic Botany 29(3): 654–669.

Hall, J.C. (2008). Systematics of Capparaceae and Cleomaceae: an evaluation of the generic delimitations of Capparis and Cleome using plastid DNA sequence data. Botany 86(7): 682–696.

Haudry, A., Platts, A.E., Vello, E., Hoen, D.R., Leclercq, M., Williamson, R.J., Forczek, E., Joly-Lopez, Z., Steffen, J.G., Hazzouri, K.M. and Dewar, K., Stinchcombe, J.R., Schoen, D.J., Wang, X., Schmutz, J., Town, C.D., Edger, P.P., Pires, J.C., Schumaker, K.S., Jarvis, D.E., Mandáková, T., Lysak, M.A., van den Bergh, E., Schranz, M.E., Harrison, P.M., Moses, A.M., Bureau, T.E., Wright, S.I., and Blanchette, M. (2013). An atlas of over 90,000 conserved noncoding sequences provides insight into crucifer regulatory regions. Nature genetics, 45(8), p.891.

Huang C.H., Sun, R., Hu, Y., Zeng, L., Zhang, N., Cai, L., Zhang, Q., Koch, M.A., Al-Shehbaz, I., Edger, P. P., and Pires, J.C. (2016). Resolution of Brassicaceae phylogeny using nuclear genes uncovers nested radiations and supports convergent morphological evolution. Molecular Biology and Evolution 33(2): 394–412.

Kagale, S., Robinson, S.J., Nixon, J., Xiao, R., Huebert, T., Condie, J., Kessler, D., Clarke, W.E., Edger, P.P., Links, M.G., Sharpe, A.G., and Parkin, I.A.P. (2014). Polyploid evolution of the Brassicaceae during the Cenozoic era. The Plant Cell, 26(7), pp.2777–2791.

Kocot, K.M., Citarella, M.R., Moroz, L.L., and Halanych, K.M. (2013). PhyloTreePruner: a phylogenetic tree-based approach for selection of orthologous sequences for phylogenomics. Evolutionary Bioinformatics 9: EBO–S12813.

Katoh, K., Misawa, K., Kuma, K.I., and Miyata, T. (2002). MAFFT: a novel method for rapid multiple sequence alignment based on fast Fourier transform. Nucleic acids research 30(14): 3059–3066.

Kearse, M., Moir, R., Wilson, A., Stones-Havas, S., Cheung, M., Sturrock, S., Buxton, S., Cooper, A., Markowitz, S., Duran, C. and Thierer, T., Ashton, B., Meintjes, P., and Drummond, A. (2012). Geneious Basic: an integrated and extendable desktop software platform for the organization and analysis of sequence data. Bioinformatics, 28(12), pp.1647–1649.

Kliebenstein, D.J., Lambrix, V.M., Reichelt, M., Gershenzon, J., and Mitchell-Olds, T. (2001). Gene duplication in the diversification of secondary metabolism: tandem 2-oxoglutarate–dependent dioxygenases control glucosinolate biosynthesis in Arabidopsis. The Plant Cell, 13(3), 681–693.

Langmead, B., and Salzberg, S.L. (2012). Fast gapped-read alignment with Bowtie2. Nature Methods 9(4): 357.

Li, Z., Tiley, G.P., Rundell, R.J., and Barker, M.S. (2019). Reply to Nakatani and McLysaght: Analyzing deep duplication events. Proceedings of the National Academy of Sciences 116(6): 1819–1820.

Li, Z, and Barker, M.S. (2019) Inferring putative ancient whole genome duplications in the 1000 Plants (1KP) initiative: access to gene family phylogenies and age distributions. bioRxiv 735076; doi: https://doi.org/10.1101/735076

Li, Z., Tiley, G.P., Galuska, S.R., Reardon, C.R., Kidder, T.I., Rundell, R.J., and Barker, M.S. (2018). Multiple large-scale gene and genome duplications during the evolution of hexapods. Proceedings of the National Academy of Sciences, 115(18), 4713–4718.

Lysak, M.A. (2018). Brassicales: an update on chromosomal evolution and ancient polyploidy. Plant Systematics and Evolution 1–6.

Lysak, M.A., Koch, M.A., Pecinka, A., and Schubert, I. (2005). Chromosome triplication found across the tribe Brassiceae. Genome Research, 15(4): 516–525.

Magallon, S., Crane, P.R., and Herendeen, P.S. (1999). Phylogenetic pattern, diversity, and diversification of eudicots. Annals of the Missouri Botanical Garden 297–372.

Mandakova, T., Li, Z., Barker, M. S., and Lysak, M. A. (2017). Diverse genome organization following 13 independent mesopolyploid events in Brassicaceae contrasts with convergent patterns of gene retention. The Plant Journal, 91(1), 3–21.

Martín-Bravo, S., Meimberg, H., Luceño, M., Märkl, W., Valcárcel, V., Bräuchler, C., Vargas, P., and Heubl, G. (2007). Molecular systematics and biogeography of Resedaceae based on ITS and trnL-F sequences. Molecular Phylogenetics and Evolution, 44(3), 1105–1120.

Martín-Bravo, S., Vargas, P., and Luceño, M. (2009). Is Oligomeris (Resedaceae) indigenous to North America? Molecular evidence for a natural colonization from the Old World. American Journal of Botany, 96(2), pp.507–518.

McKain, M.R., Wickett, N., Zhang, Y., Ayyampalayam, S., McCombie, W.R., Chase, M.W., Pires, J.C., de Pamphilis, C.W., and Leebens-Mack, J. (2012). Phylogenomic analysis of transcriptome data elucidates co-occurrence of a paleopolyploid event and the origin of bimodal karyotypes in Agavoideae (Asparagaceae). American journal of botany, 99(2), pp.397–406.

McKain M.R., and Wilson, M. (2017). Fast-Plast: rapid de novo assembly and finishing for whole chloroplast genomes. Version 1.2.6

McKain, M.R., Tang, H., McNeal, J.R., Ayyampalayam, S., Davis, J.I., dePamphilis, C.W., Givnish, T.J., Pires, J.C., Stevenson, D.W., and Leebens-Mack, J.H. (2016). A phylogenomic assessment of ancient polyploidy and genome evolution across the Poales. Genome biology and evolution, 8(4), pp.1150–1164.

Ming, R., Hou, S., Feng, Y., Yu, Q., Dionne-Laporte, A., Saw, J.H., Senin, P., Wang, W., Ly, B.V., Lewis, K.L. and Salzberg, S.L., Feng, L., Jones, M. R., Skelton, R.L., Murray, J.E., Chen, C., Qian, W., Shen, J., Du, P., Eustice, M., Tong, E., Tang, H., Lyons, E., Paull, R.E., Michael, T.P., Wall, K., Rice, D.W., Albert, H., Wang, M., Zhu, Y.J., Schatz, M., Nagarajan, N., Acob, R.A., Guan, P., Blas, A., Wai, C.M., Ackerman, C.M., Ren, Y., Liu, C., Wang, J., Wang, J., Na, J., Shakirov, E.V., Haas, B., Thimmapuram, J., Nelson, D., Wang, X., Bowers, J.E., Gschwend, A.R., Delcher, A.L., Singh, R., Suzuki, J.Y., Tripathi, S., Neupane, K., Wei, H., Irikura, B., Paidi, M., Jiang, N., Zhang, W., Presting, G., Windsor, A., Navajas-Pérez, R., Torres, M.J., Feltus, F.A., Porter, B., Li, Y., Burroughs, A.M., Luo, M., Liu, L., Christopher, D.A., Mount, S.M., Moore, P.H., Sugimura, T., Jiang, J., Schuler, M.A., Friedman, V., Mitchell-Olds, T., Shippen, D.E., dePamphilis, C.W., Palmer, J.D., Freeling, M., Paterson, A.H., Gonsalves, D., Wang, L., and Alam, M. (2008). The draft genome of the transgenic tropical fruit tree papaya (*Carica papaya* Linnaeus). Nature, 452(7190), p.991.

Mithen, R., Bennett, R., and Marquez, J. (2010). Glucosinolate biochemical diversity and innovation in the Brassicales. Phytochemistry 71(17-18): 2074–2086.

Naik, P. A., Shi, P., and Tsai, C. L. (2007). Extending the Akaike information criterion to mixture regression models. Journal of the American Statistical Association 102(477): 244–254.

Nakatani, Y., and McLysaght, A. (2019). Macrosynteny analysis shows the absence of ancient whole-genome duplication in lepidopteran insects. Proceedings of the National Academy of Sciences 116(6): 1816–1818.

Nikolov, L.A., Shushkov, P., Nevado, B., Gan, X., Al-Shehbaz, I.A., Filatov, D., Bailey, C.D., and Tsiantis, M. (2019). Resolving the backbone of the Brassicaceae phylogeny for investigating trait diversity. New Phytologist, 222(3), pp.1638–1651.

Patchell, M. J., Roalson, E. H., and Hall, J. C. (2014). Resolved phylogeny of Cleomaceae based on all three genomes. Taxon, 63(2), 315–328.

R Core Team. (2018). R: A language and environment for statistical computing. R Foundation for Statistical Computing, Vienna, Austria. URL https://www.R-project.org/.

Ratzka, A., Vogel, H., Kliebenstein, D.J., Mitchell-Olds, T., and Kroymann, J. (2002). Disarming the mustard oil bomb. Proceedings of the National Academy of Sciences, 99(17), 11223–11228.

Rodman, J. E., Soltis, P. S., Soltis, D. E., Sytsma, K. J., and Karol, K. G. (1998). Parallel evolution of glucosinolate biosynthesis inferred from congruent nuclear and plastid gene phylogenies. American Journal of Botany, 85(7), 997–1006.

Schlüter, U., Bräutigam, A., Gowik, U., Melzer, M., Christin, P.A., Kurz, S., Mettler-Altmann, T., and Weber, A.P. (2016). Photosynthesis in C3–C4 intermediate Moricandia species. Journal of Experimental Botany, 68(2), 191–206.

Schranz M. E. and Mitchell-Olds, T. (2006). Independent ancient polyploidy events in the sister families Brassicaceae and Cleomaceae. Plant Cell 18(5):1152–1165

Simão, F. A., Waterhouse, R. M., Ioannidis, P., Kriventseva, E.V., and Zdobnov, E. M. (2015). BUSCO: assessing genome assembly and annotation completeness with single-copy orthologs. Bioinformatics 31(19): 3210–3212.

Smith S.A. and Dunn, C.W. (2008). Phyutility: a phyloinformatics tool for trees, alignments, and molecular data. Bioinformatics 24: 715–716.

Stamatakis, A. (2014). RAxML version 8: a tool for phylogenetic analysis and post-analysis of large phylogenies. Bioinformatics 30(9): 1312–1313.

Tamboli, A. S., Yadav, P. B., Gothe, A. A., Yadav, S. R., and Govindwar, S. P. (2018). Molecular phylogeny and genetic diversity of genus Capparis (Capparaceae) based on plastid DNA sequences and ISSR markers. Plant systematics and evolution, 304(2), 205–217.

Tian, Y., Zeng, Y., Zhang, J., Yang, C., Yan, L., Wang, X., Shi, C., Xie, J., Dai, T., Peng, L. and Huan, Y.Z., Xu, A., Huang, Y., Zhang, J., Ma, X., Dong, Y., Hao, S., and Sheng, J. (2015). High quality reference genome of drumstick tree (*Moringa oleifera* Lam.), a potential perennial crop. Science China Life Sciences, 58(7), pp.627–638.

Tiley, G. P., Barker, M. S., and Burleigh, J. G. (2018). Assessing the performance of Ks plots for detecting ancient Whole-genome duplications. Genome Biology and Evolution 10(11): 2882–2898.

van den Bergh E., Külahoglu, C., Bräutigam, A., Hibberd, J. M., Weber, A. P. M., Zhu, X. G., and Schranz, M. E. (2014). Gene and genome duplications and the origin of C4 photosynthesis: Birth of a trait in the Cleomaceae. Current Plant Biology 1: 2–9.

Vision, T. J., Brown, D. G., and Tanksley, S. D. (2000). The origins of genomic duplications in *Arabidopsis*. Science 290(5499): 2114–2117.

Waterhouse, R.M., Seppey, M., Simão, F.A., Manni, M., Ioannidis, P., Klioutchnikov, G., Kriventseva, E.V., and Zdobnov, E.M. (2017). BUSCO applications from quality assessments to gene prediction and phylogenomics. Molecular biology and evolution, 35(3), pp.543–548.

Washburn, J.D., Schnable, J.C., Conant, G.C., Brutnell, T.P., Shao, Y., Zhang, Y., Ludwig, M., Davidse, G.. and Pires, J.C. (2017). Genome-Guided Phylo-Transcriptomic Methods and the Nuclear Phylogenetic Tree of the Paniceae Grasses. Scientific reports, 7(1), p.13528.

Yang, Y., Moore, M.J., Brockington, S.F., Soltis, D.E., Wong, G.K.S., Carpenter, E.J., Zhang, Y., Chen, L., Yan, Z., Xie, Y. and Sage, R.F., Covshoff, S., Hibberd, J.M., Nelson, M.N., and Smith, S.A. (2015). Dissecting molecular evolution in the highly diverse plant clade Caryophyllales using transcriptome sequencing. Molecular Biology and Evolution, 32(8), pp.2001–2014.

Zhang, C., Rabiee, M., Sayyari, E., and Mirarab, S. (2018). ASTRAL-III: polynomial time species tree reconstruction from partially resolved gene trees. BMC bioinformatics 19(6): 153.

Züst, T., Mirzaei, M., and Jander, G. (2018). Erysimum cheiranthoides, an ecological research system with potential as a genetic and genomic model for studying cardiac glycoside biosynthesis. Phytochemistry reviews, 17(6), 1239–1251.

Zwaenepoel, A., Li, Z., Lohaus, R., and Van de Peer, Y. (2019). Finding evidence for Whole-genome duplications: a reappraisal. Molecular Plant 12(2): 133–136.

Zwaenepoel, A., and Van de Peer, Y. (2019). Ancient Whole-genome duplications and the evolution of the gene duplication and loss rate. bioRxiv 556076.

