## Supplemental Figures and Tables for "Phylogeny and Multiple Independent Whole-Genome Duplication Events in the Brassicales"

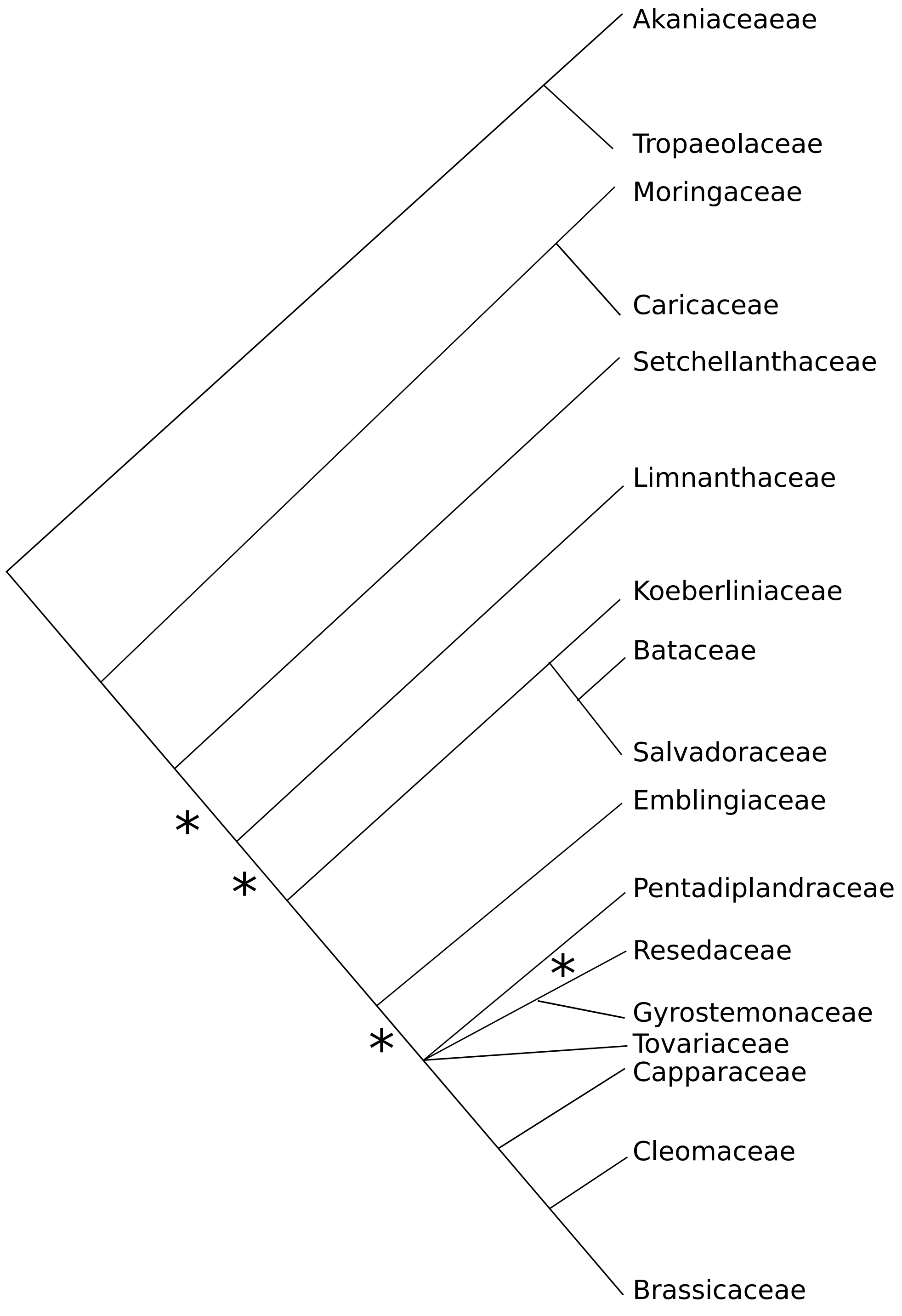

Supp. Figure 1. Current understanding of the relationships between the 17 families of the Brassicales (APG IV). * indicates branch support between 50-80%, all other branches have greater than 80% support.

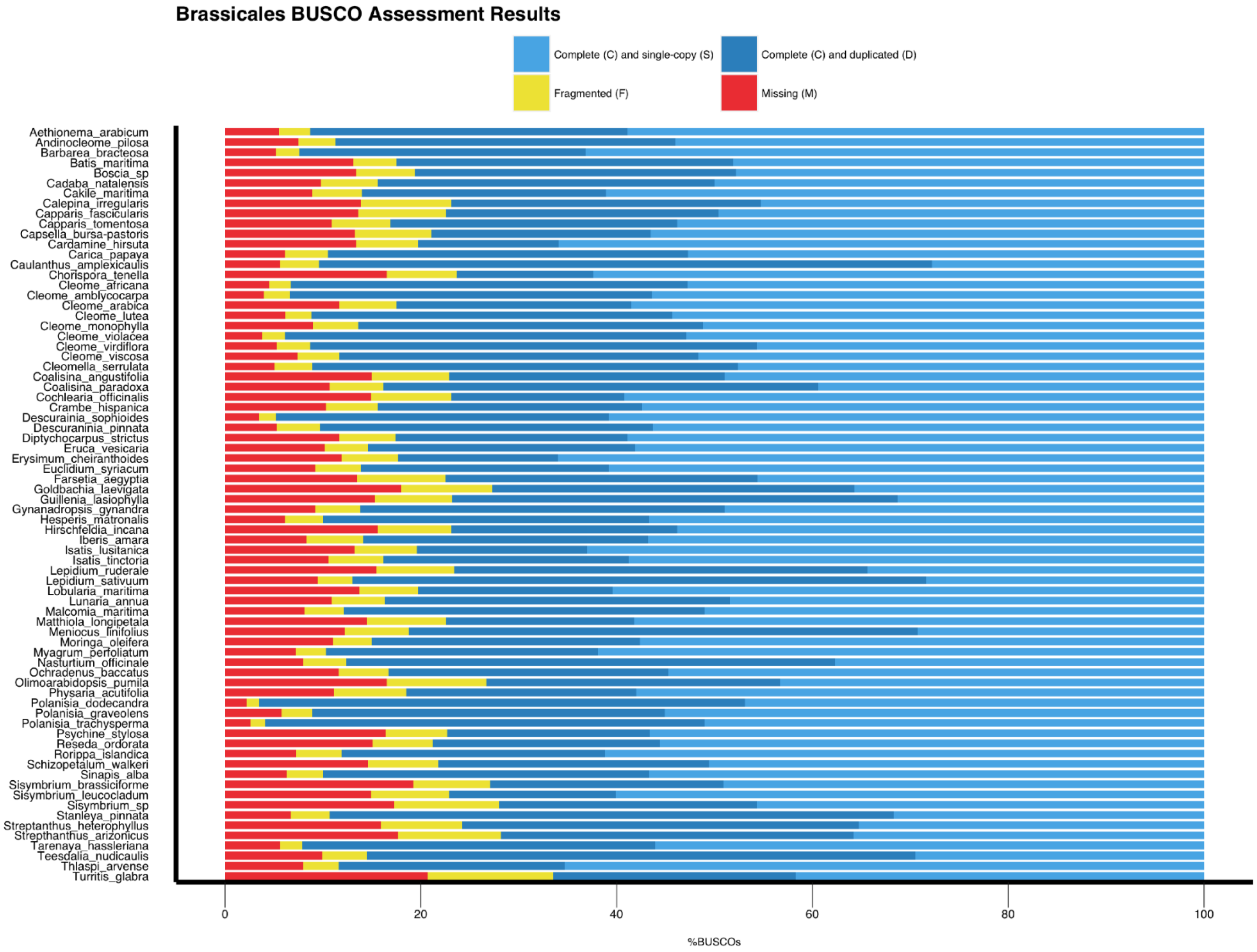

Supp. Figure 2. BUSCO analysis of de novo transcriptomes. Legend indicates the percent of genes that are complete and single copy (light blue), complete and duplicate (dark blue), fragmented (yellow), and missing (red) in de novo transcriptomes.

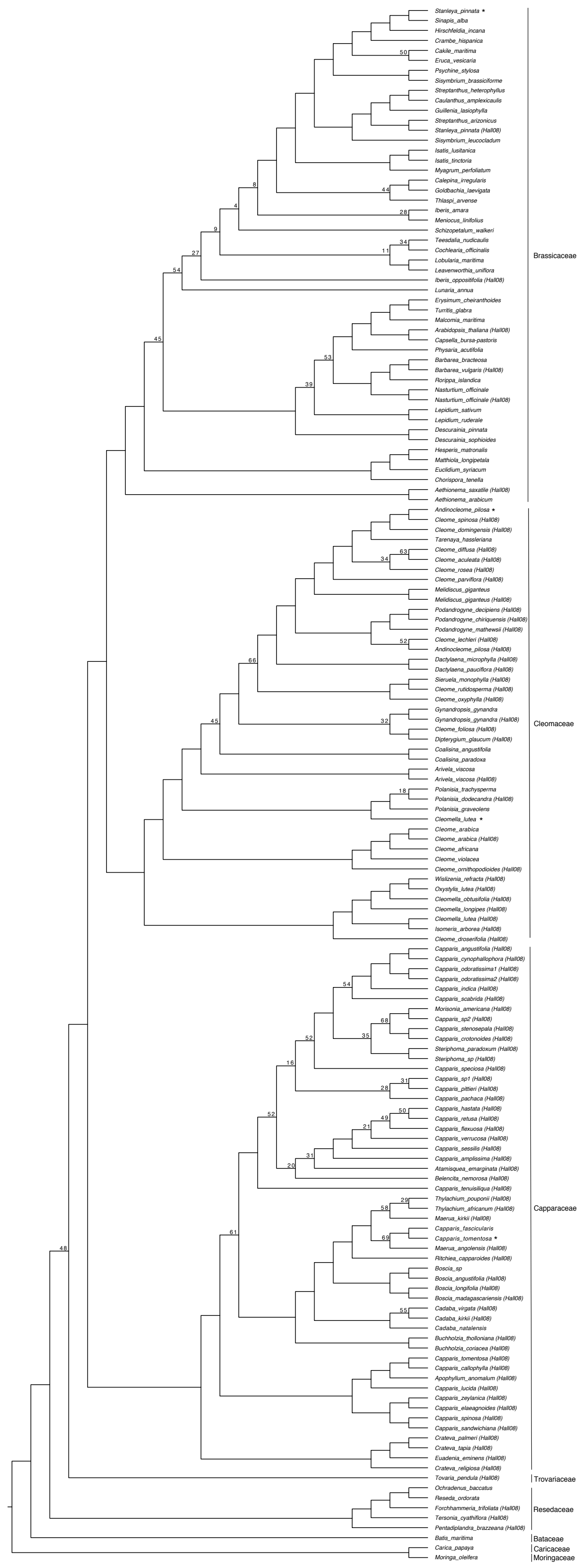

Supp. Figure 3. Maximum likelihood phylogeny of the Brassicales using two chloroplast genes, MatK and NdhF. Support values are indicated if below 70 percent bootstrap support. * next to taxa indicate those whose placement are not sister with samples from Hall (2008).

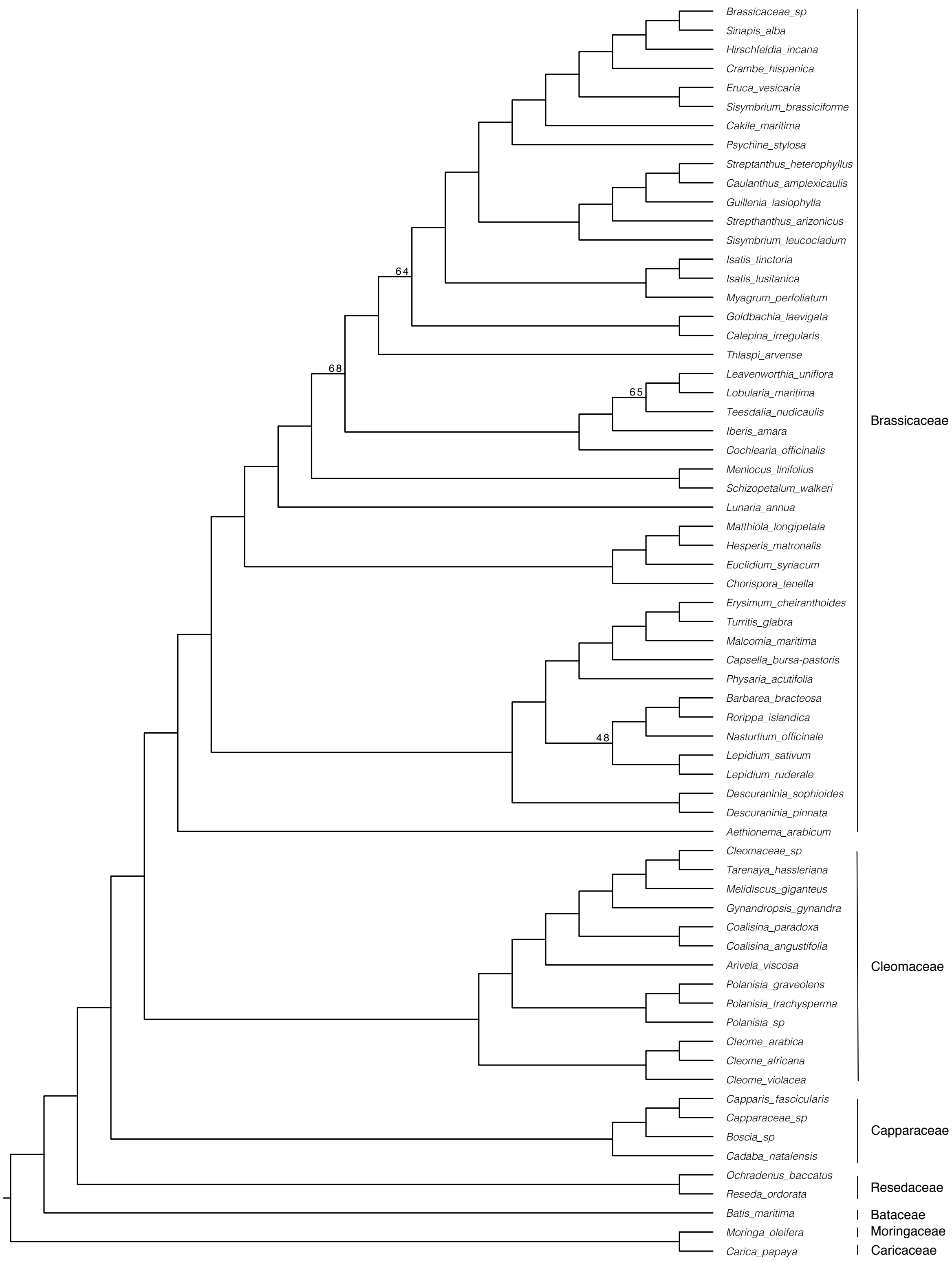

Supp. Figure 4. Maximum likelihood whole-chloroplast phylogeny of the Brassicales. Support values are indicated if below 70 percent bootstrap support.

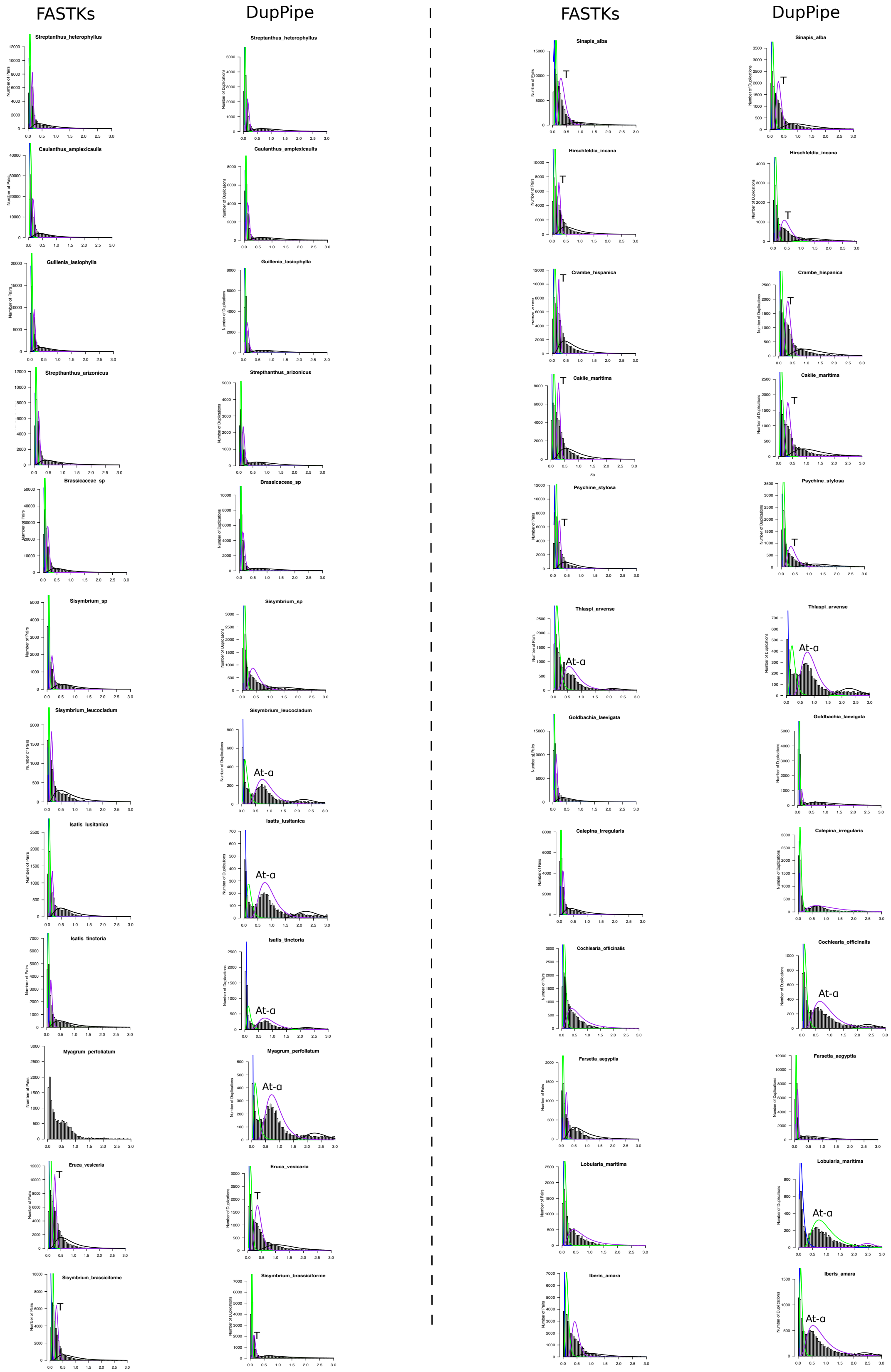

FASTKs
DupPipe
FASTKs
DupPipe
Streptanthus_heterophyllus
Caulanthus_amplexicaulis
Guillenia_lasiophylla
Strepthanthus_arizonicus
Brassicaceae_sp
Sisymbrium_sp
Sisymbrium_leucocladum
Isatis_lusitanica
Isatis_tinctoria
Myagrum_perfoliatum
Eruca_vesicaria
Sisymbrium_brassiciforme
Sinapis_alba
Hirschfeldia_incana
Crambe_hispanica
Cakile_maritima
Psychine_stylosa
Thlaspi_arvense
Goldbachia_laevigata
Calepina_irregularis
Cochlearia_officinalis
Farsetia_aegyptia
Lobularia_maritima
Iberis_amara
T
At-α
Number of Pairs
Number of Duplications
Ks

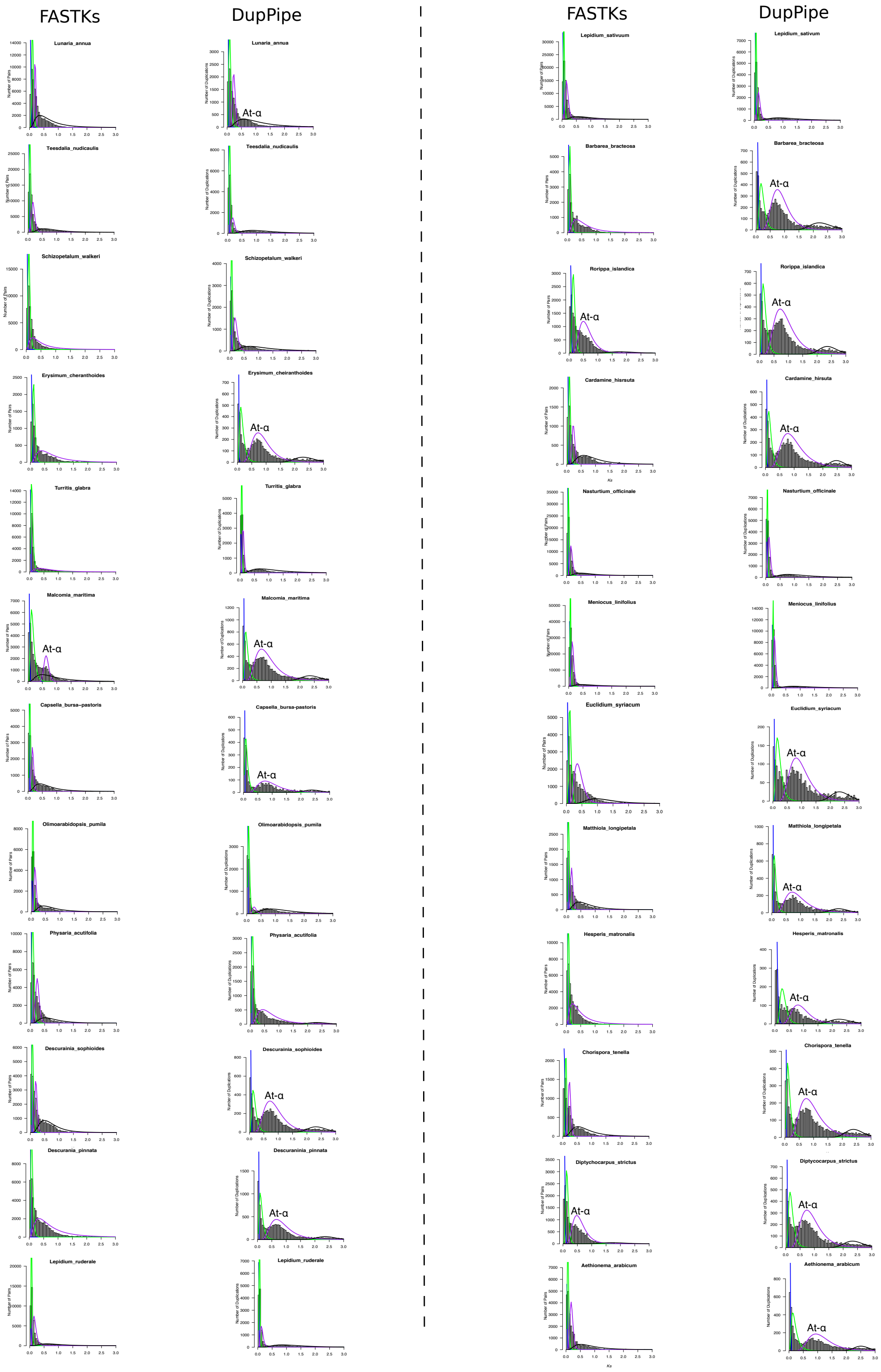

Supp. Figure 5. Ks plots of the Brassicaceae using both FASTKs (McKain et al. 2016) and DupPipe (Barker et al. 2010). Whole-genome duplication events, At-ɑ and the Brassiceae triplication (T) event are noted above corresponding peaks.

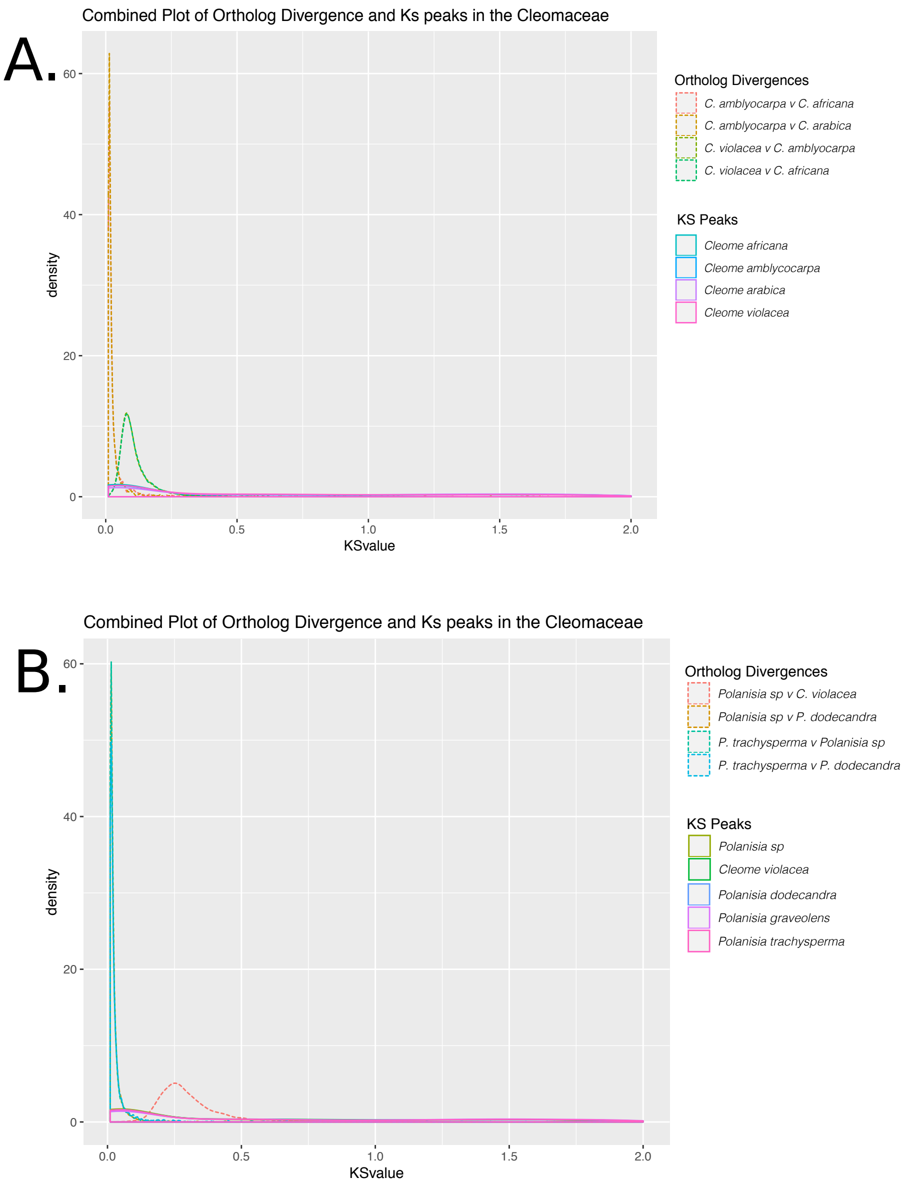

Supp. Figure 6. Ortholog divergences and Ks peaks of the Cleomaceae. **(A)** Ortholog divergences between *C. amblyocarpa* and *C. africana, C. arabica,* and *C. violacea* and between *C. violacea* and *C. africana* to test placement of potential novel WGD event. **(B)** Ortholog divergences between *Polanisia sp.* and *C. violacea, Polanisia sp.* and *P. dodecandria, P. trachysperma* and *Polanisia sp.*, and between *P. trachysperma* and *P. dodecandria* to test for placement of the second potential novel WGD event.

A.

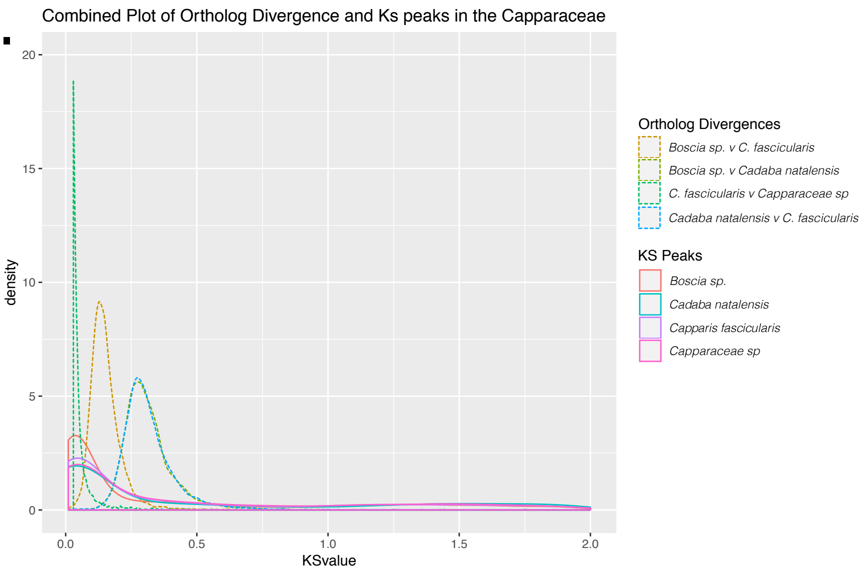

B.

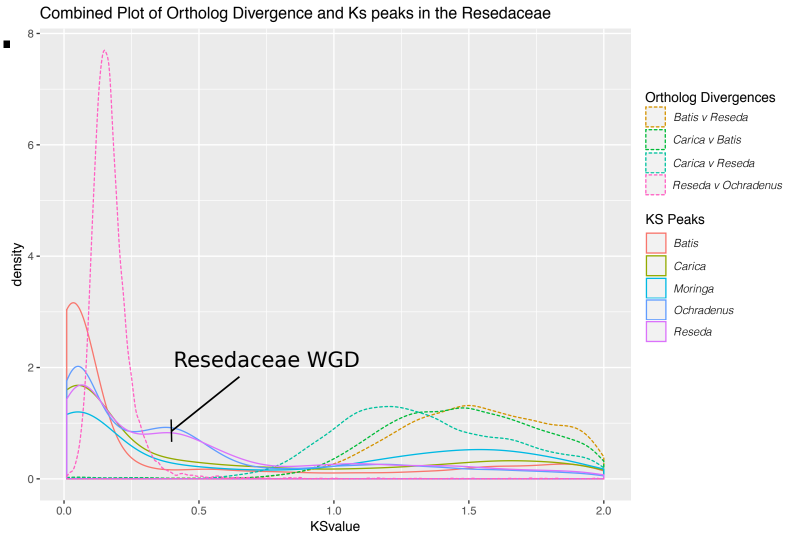

Supp. Figure 7. Ortholog divergences and Ks peaks of the **(A)** Capparaceae and **(B)** Resedaceae + Outgroups. Proposed Resedaceae whole-genome duplication event indicated.

Supp. Table 1

| Family | Genus | Species | Seed Accession | Collection | Transcriptome | Chloroplast SSC | Chloroplast LSC | Chloroplast IR | Fast-Plast Subsample | SRA RNA-seq # | SRA GSS # |
|---|---|---|---|---|---|---|---|---|---|---|---|
| **Bataceae** | *Batis* | *maritima* | *NA* | Pat Edger | X | X | X | X | 3500000 | | |
| **Brassicaceae** | *Aethionema* | *arabicum* | 1010 | Eric Schranz | X | X | X | X | | | |
| **Brassicaceae** | *Barbarea* | *bracteosa* | 6203-84 | Gomez Campo | X | X | X | X | | | |
| **Brassicaceae** | *Brassicaceae* | *sp* | Colorado diploid | Chris Pires | X | X | X | X | | | |
| **Brassicaceae** | *Cakile* | *maritima* | B2005-875 | Marcus Koch | X | X | X | X | | | |
| **Brassicaceae** | *Calepina* | *irregularis* | 2158-72 | Gomez Campo | X | X | X | X | | | |
| **Brassicaceae** | *Capsella* | *bursa-pastoris* | 2006-956 | Marcus Koch | X | X | X | X | | | |
| **Brassicaceae** | *Cardamine* | *hirsuta* | 2005-701 | Marcus Koch | X | X | | | | | |
| **Brassicaceae** | *Caulanthus* | *amplexicaulis* | *NA* | Alan Pepper | X | X | X | X | | | |
| **Brassicaceae** | *Chorispora* | *tenella* | 2006-629 | Marcus Koch | X | X | X | X | 1000000 | | |
| **Brassicaceae** | *Cochlearia* | *officinalis* | 1910-84 | Gomez Campo | X | X | X | X | | | |
| **Brassicaceae** | *Crambe* | *hispanica* | PI388853 | USDA | X | X | X | X | | | |
| **Brassicaceae** | *Descurainia* | *pinnata* | B2004-26 | Marcus Koch | X | X | X | X | | | |
| **Brassicaceae** | *Descurainia* | *sophioides* | BarbGH | Chris Pires | X | X | X | X | | | |
| **Brassicaceae** | *Diptycocarpus* | *strictus* | 05-0397-10-00 | Klaus Mummenhoff | X | | | | | | |
| **Brassicaceae** | *Eruca* | *vesicaria* | PI633218 | USDA | X | X | X | X | 1000000 | | |
| **Brassicaceae** | *Erysimum* | *cheiranthoides* | 03-0030-10-00 | Klaus Mummenhoff | X | X | X | X | | | |
| **Brassicaceae** | *Euclidium* | *syriacum* | 0587-68 | Gomez Campo | X | X | X | X | | | |
| **Brassicaceae** | *Farsetia* | *aegyptia* | 96-001-10-00 | Klaus Mummenhoff | X | | X | | | | |
| **Brassicaceae** | *Goldbachia* | *laevigata* | *NA* | Martin Lysak | X | X | X | X | | | |
| **Brassicaceae** | *Guillenia* | *lasiophylla* | Costal type | Chris Pires | X | X | X | X | | | |
| **Brassicaceae** | *Hesperis* | *matronalis* | 0039-68 | Gomez Campo | X | X | X | X | 1000000 | | |
| **Brassicaceae** | *Hirschfeldia* | *incana* | 2024-71 | Gomez Campo | X | X | X | X | | | |
| **Brassicaceae** | *Iberis* | *amara* | 2005-238 | Marcus Koch | X | X | X | X | | | |
| **Brassicaceae** | *Isatis* | *lusitanica* | 01-0035-10-00 | Klaus Mummenhoff | X | X | X | X | | | |
| **Brassicaceae** | *Isatis* | *tinctoria* | B2009-0378 | Marcus Koch | X | X | X | X | | | |
| **Brassicaceae** | *Leavenworthia* | *uniflora* | 2004-557 | Marcus Koch | | X | X | X | | | |

| | | | | | | | | | |
|---|---|---|---|---|---|---|---|---|---|
| **Brassicaceae** | *Lepidium* | *ruderale* | 05-0190-10-00 | Klaus Mummenhoff | X | X | X | X | |
| **Brassicaceae** | *Lepidium* | *sativum* | 2003-169 | Marcus Koch | X | X | X | X | |
| **Brassicaceae** | *Lobularia* | *maritima* | B2009-0390 | Marcus Koch | X | X | X | X | |
| **Brassicaceae** | *Lunaria* | *annua* | 2006-705 | Marcus Koch | X | X | X | X | |
| **Brassicaceae** | *Malcomia* | *maritima* | 2008-777 | Marcus Koch | X | X | X | X | 2500000 |
| **Brassicaceae** | *Matthiola* | *longipetala* | 1351-70 | Gomez Campo | X | X | X | X | |
| **Brassicaceae** | *Meniocus* | *linifolius* | 0893-68 | Gomez Campo | X | X | X | X | 1000000 |
| **Brassicaceae** | *Myagrum* | *perfoliatum* | 2006-432 | Marcus Koch | X | X | X | X | |
| **Brassicaceae** | *Nasturtium* | *officinale* | AMC7.20.15 | Chris Pires | X | X | X | X | |
| **Brassicaceae** | *Olimoarabidopsis* | *pumila* | BarbGH | Chris Pires | X | | | | |
| **Brassicaceae** | *Physaria* | *acutifolia* | 01201.13 ALPLAINS 50 | Chris Pires | X | X | X | X | |
| **Brassicaceae** | *Psychine* | *stylosa* | B2004-78 | Marcus Koch | X | X | X | X | |
| **Brassicaceae** | *Rorippa* | *islandica* | 2006-89 | Gomez Campo | X | X | X | X | |
| **Brassicaceae** | *Schizopetalum* | *walkeri* | *NA* | Martin Lysak | X | X | X | X | |
| **Brassicaceae** | *Sinapis* | *alba* | 0560-79 | Gomez Campo | X | X | X | X | 1000000 |
| **Brassicaceae** | *Sisymbrium* | *brassiciforme* | 04-0372-10-00 | Klaus Mummenhoff | X | X | X | X | |
| **Brassicaceae** | *Sisymbrium* | *leucocladum* | 3760-75 | Gomez Campo | X | X | X | X | |
| **Brassicaceae** | *Sisymbrium* | *sp* | 2004-557 | Marcus Koch | X | | | | |
| **Brassicaceae** | *Streptanthus* | *arizonicus* | 02-0095-00-00 | Klaus Mummenhoff | X | X | X | X | |
| **Brassicaceae** | *Streptanthus* | *heterophyllus* | 1730-71 | Gomez Campo | X | X | X | X | |
| **Brassicaceae** | *Teesdalia* | *nudicaulis* | 2219-73 | Gomez Campo | X | X | X | X | |
| **Brassicaceae** | *Thlaspi* | *arvense* | 1211-67 | Gomez Campo | X | X | X | X | |
| **Brassicaceae** | *Turritis* | *glabra* | 04-0314-10-00 | Klaus Mummenhoff | X | X | X | X | 4500000 |
| **Capparaceae** | *Boscia* | *sp* | SHN 12-075 | Silverhill seeds and books | X | X | X | X | |
| **Capparaceae** | *Cadaba* | *natalensis* | SHN 12-076 | Silverhill seeds and books | X | X | X | X | |
| **Capparaceae** | *Capparaceae* | *sp* | SHN 12-079 | Silverhill seeds and books | X | X | X | X | |
| **Capparaceae** | *Capparis* | *fascicularis* | SHN 12-078 | Silverhill seeds and books | X | X | X | X | |
| **Cariaceae** | *Carica* | *papaya* | *NA* | Chris Pires | X | X | X | X | |

| | | | | | | | | | |
|---|---|---|---|---|---|---|---|---|---|
| **Cleomaceae** | *Arivela* | *viscosa* | KEW 104126 | Kew Gardens | X | X | X | X | |
| **Cleomaceae** | *Cleomaceae* | *sp* | KEW 36588 | Kew Gardens | X | X | X | X | |
| **Cleomaceae** | *Cleome* | *africana* | *NA* | Chris Pires | X | X | X | X | |
| **Cleomaceae** | *Cleome* | *amblycocarpa* | 151485 | Jocelyn Hall | X | | | | |
| **Cleomaceae** | *Cleome* | *arabica* | *NA* | Jocelyn Hall | X | X | X | X | |
| **Cleomaceae** | *Cleome* | *violacea* | *NA* | Jocelyn Hall | X | X | X | X | |
| **Cleomaceae** | *Cleomella* | *serrulata* | *NA* | Jocelyn Hall | X | | | | |
| **Cleomaceae** | *Coalisina* | *angustifolia* | KEW 82633 | Kew Gardens | X | X | X | X | 2000000 |
| **Cleomaceae** | *Coalisina* | *paradoxa* | KEW 109051 | Kew Gardens | X | X | X | X | 1000000 |
| **Cleomaceae** | *Gynandropsis* | *gynandra* | *NA* | Jocelyn Hall | X | X | X | X | |
| **Cleomaceae** | *Melidiscus* | *giganteus* | *NA* | Jocelyn Hall | X | X | X | X | |
| **Cleomaceae** | *Polanisia* | *dodecandra* | 2006-372 | Marcus Koch | X | X | | | |
| **Cleomaceae** | *Polanisia* | *graveolens* | 2004 -157 | Marcus Koch | X | X | X | X | |
| **Cleomaceae** | *Polanisia* | *sp* | KEW 93745 | Kew Gardens | X | X | X | X | |
| **Cleomaceae** | *Polanisia* | *trachysperma* | 2006-365 | Marcus Koch | X | X | X | X | 1600000 |
| **Cleomaceae** | *Sieruela* | *monophylla* | KEW 413163 | Kew Gardens | X | | | | |
| **Cleomaceae** | *Tarenaya* | *hassleriana* | KEW 48516 | Kew Gardens | X | X | X | X | |
| **Moringaceae** | *Moringa* | *oleifera* | *NA* | Chris Pires | X | X | X | X | 1000000 |
| **Resadaceae** | *Ochradenus* | *baccatus* | DMJ0324 | Chris Pires | X | X | X | X | |
| **Resadaceae** | *Reseda* | *ordorata* | *NA* | Larry Chandler | X | X | X | X | 1000000 |

| Group | # of genes assigned to groups | % assigned | OrthoGroups | G50 | O50 | # of groups with all species | Single-copy OrthoGroups | Taxon Occupancy (80%) | Alignment Quality (40% gaps) | # of Trees after first round of RAxML | Tree Pruning (cutoff #) | Final tree # for species tree inference |
|---|---|---|---|---|---|---|---|---|---|---|---|---|
| **Brassicales** | 3516602 | 96.3 | 47600 | 252 | 4630 | 6444 | 0 | 59/74 = 10968 | 2663 | 2663 | 1284 (10) | 1284 |
| **Brassicaceae** | 2224281 | 96.7 | 39809 | 135 | 5160 | 7822 | 0 | 39/48 = 12129 | 5587 | 5587 | 2110 (10) | 2110 |
| **Capparaceae** | 143114 | 79.4 | 25179 | 5 | 9600 | 12350 | 3228 | 3/4 = 17571 | 15988 | 14338 | 13328 (3) | 10214 |
| **Cleomaceae** | 883407 | 91.9 | 42218 | 37 | 7662 | 9706 | 55 | 13/17 = 14107 | 10732 | 10732 | 3626 (10) | 3626 |
| **RBMC** | 165057 | 78.7 | 21372 | 8 | 7447 | 9098 | 881 | 4/5 = 11756 | 10038 | 10038 | 8476 (4) | 8476 |

Supp. Table 3

| Family | Genus | Species | RNA Extraction Kit | RNA Library Prep | RNA Seq Machine | RNA Read Length | RNA Raw Reads | DNA Extraction Kit | DNA Library Prep | DNA Seq Machine | DNA Read Length | DNA Raw Reads |
|---|---|---|---|---|---|---|---|---|---|---|---|---|
| **Bataceae** | *Batis* | *maritima* | ThermoFisher PureLink RNA Mini Kit | TruSeq | NextSeq | 2X75 | 16565629 | Qiagen DNeasy | TruSeq | NextSeq | 2X150 | 8309055 |
| **Brassicaceae** | *Aethionema* | *arabicum* | ThermoFisher PureLink RNA Mini Kit | TruSeq | NextSeq | 2X75 | 55433848 | Qiagen DNeasy | TruSeq | NextSeq | 2X150 | 8757856 |
| **Brassicaceae** | *Barbarea* | *bracteosa* | ThermoFisher PureLink RNA Mini Kit | TruSeq | NextSeq | 2X75 | 53269435 | Qiagen DNeasy | TruSeq | NextSeq | 2X150 | 8979330 |
| **Brassicaceae** | *Brassicaceae* | *sp* | ThermoFisher PureLink RNA Mini Kit | TruSeq | HiSeq | 2X100 | 40618374 | Qiagen DNeasy | TruSeq | NextSeq | 2X150 | 7179797 |
| **Brassicaceae** | *Cakile* | *maritima* | ThermoFisher PureLink RNA Mini Kit | TruSeq | HiSeq | 2X250 | 5555024 | Qiagen DNeasy | TruSeq | NextSeq | 2X150 | 10054961 |
| **Brassicaceae** | *Calepina* | *irregularis* | ThermoFisher PureLink RNA Mini Kit | TruSeq | NextSeq | 2X75 | 16367748 | Qiagen DNeasy | TruSeq | NextSeq | 2X150 | 9472593 |
| **Brassicaceae** | *Capsella* | *bursa-pastoris* | ThermoFisher PureLink RNA Mini Kit | TruSeq | NextSeq | 2X75 | 23770081 | Qiagen DNeasy | TruSeq | NextSeq | 2X150 | 9962200 |
| **Brassicaceae** | *Cardamine* | *hirsuta* | ThermoFisher PureLink RNA Mini Kit | TruSeq | NextSeq | 2X75 | 23276338 | Qiagen DNeasy | TruSeq | NextSeq | 2X150 | 7163619 |
| **Brassicaceae** | *Caulanthus* | *amplexicaulis* | ThermoFisher PureLink RNA Mini Kit | TruSeq | HiSeq | 2X100 | 30561937 | Qiagen DNeasy | TruSeq | NextSeq | 2X150 | 8638165 |
| **Brassicaceae** | *Chorispora* | *tenella* | ThermoFisher PureLink RNA Mini Kit | TruSeq | NextSeq | 2X75 | 19065280 | Qiagen DNeasy | TruSeq | NextSeq | 2X150 | 10713329 |
| **Brassicaceae** | *Cochlearia* | *officinalis* | ThermoFisher PureLink RNA Mini Kit | TruSeq | NextSeq | 2X75 | 19278016 | Qiagen DNeasy | TruSeq | NextSeq | 2X150 | 9785784 |
| **Brassicaceae** | *Crambe* | *hispanica* | ThermoFisher PureLink RNA Mini Kit | TruSeq | HiSeq | 2X100 | 6677521 | Qiagen DNeasy | TruSeq | NextSeq | 2X150 | 8981402 |
| **Brassicaceae** | *Descurainia* | *pinnata* | ThermoFisher PureLink RNA Mini Kit | TruSeq | HiSeq | 2X100 | 8315473 | Qiagen DNeasy | TruSeq | NextSeq | 2X150 | 10532085 |
| **Brassicaceae** | *Descurainia* | *sophioides* | ThermoFisher PureLink RNA Mini Kit | TruSeq | HiSeq | 2X100 | 41235763 | Qiagen DNeasy | TruSeq | NextSeq | 2X150 | 7149869 |
| **Brassicaceae** | *Diptycocarpus* | *strictus* | ThermoFisher PureLink RNA Mini Kit | TruSeq | HiSeq | 2X100 | 7422199 | | | | | 7488205 |
| **Brassicaceae** | *Eruca* | *vesicaria* | ThermoFisher PureLink RNA Mini Kit | TruSeq | HiSeq | 2X100 | 7743240 | Qiagen DNeasy | TruSeq | NextSeq | 2X150 | 9696734 |
| **Brassicaceae** | *Erysimum* | *cheiranthoides* | ThermoFisher PureLink RNA Mini Kit | TruSeq | NextSeq | 2X75 | 19481894 | Qiagen DNeasy | TruSeq | NextSeq | 2X150 | 9627950 |
| **Brassicaceae** | *Euclidium* | *syriacum* | ThermoFisher PureLink RNA Mini Kit | TruSeq | HiSeq | 2X100 | 7871790 | Qiagen DNeasy | TruSeq | NextSeq | 2X150 | 8499185 |
| **Brassicaceae** | *Farsetia* | *aegyptia* | ThermoFisher PureLink RNA Mini Kit | TruSeq | NextSeq | 2X75 | 49764414 | Qiagen DNeasy | TruSeq | NextSeq | 2X150 | 8514984 |
| **Brassicaceae** | *Goldbachia* | *laevigata* | ThermoFisher PureLink RNA Mini Kit | TruSeq | NextSeq | 2X75 | 16809960 | Qiagen DNeasy | TruSeq | NextSeq | 2X150 | 9849790 |
| **Brassicaceae** | *Guillenia* | *lasiophylla* | ThermoFisher PureLink RNA Mini Kit | TruSeq | NextSeq | 2X75 | 23503171 | Qiagen DNeasy | TruSeq | NextSeq | 2X150 | 10824454 |
| **Brassicaceae** | *Hesperis* | *matronalis* | ThermoFisher PureLink RNA Mini Kit | TruSeq | NextSeq | 2X75 | 51877498 | Qiagen DNeasy | TruSeq | NextSeq | 2X150 | 7807320 |
| **Brassicaceae** | *Hirschfeldia* | *incana* | ThermoFisher PureLink RNA Mini Kit | TruSeq | NextSeq | 2X75 | 18013197 | Qiagen DNeasy | TruSeq | NextSeq | 2X150 | 11527537 |
| **Brassicaceae** | *Iberis* | *amara* | ThermoFisher PureLink RNA Mini Kit | TruSeq | HiSeq | 2X100 | 7587536 | Qiagen DNeasy | TruSeq | NextSeq | 2X150 | 6845991 |
| **Brassicaceae** | *Isatis* | *lusitanica* | ThermoFisher PureLink RNA Mini Kit | TruSeq | NextSeq | 2X75 | 22464165 | Qiagen DNeasy | TruSeq | NextSeq | 2X150 | 11695349 |

| | | | | | | | | | | | | |
|---|---|---|---|---|---|---|---|---|---|---|---|---|
| **Brassicaceae** | *Isatis* | *tinctoria* | Qiagen RNEasy | TruSeq | HiSeq | 2X100 | 20649046 | Qiagen DNeasy | TruSeq | NextSeq | 2X150 | 9446020 |
| **Brassicaceae** | *Leavenworthia* | *uniflora* | NA | NA | NA | NA | NA | Qiagen DNeasy | TruSeq | NextSeq | 2X150 | 9120176 |
| **Brassicaceae** | *Lepidium* | *ruderale* | ThermoFisher PureLink RNA Mini Kit | TruSeq | NextSeq | 2X75 | 18623760 | Qiagen DNeasy | TruSeq | NextSeq | 2X150 | 9459705 |
| **Brassicaceae** | *Lepidium* | *sativum* | ThermoFisher PureLink RNA Mini Kit | TruSeq | HiSeq | 2X100 | 7083621 | Qiagen DNeasy | TruSeq | NextSeq | 2X150 | 9451177 |
| **Brassicaceae** | *Lobularia* | *maritima* | ThermoFisher PureLink RNA Mini Kit | TruSeq | NextSeq | 2X75 | 21143830 | Qiagen DNeasy | TruSeq | NextSeq | 2X150 | 9578197 |
| **Brassicaceae** | *Lunaria* | *annua* | ThermoFisher PureLink RNA Mini Kit | TruSeq | HiSeq | 2X100 | 6184646 | Qiagen DNeasy | TruSeq | NextSeq | 2X150 | 7316600 |
| **Brassicaceae** | *Malcomia* | *maritima* | ThermoFisher PureLink RNA Mini Kit | TruSeq | HiSeq | 2X100 | 7408880 | Qiagen DNeasy | TruSeq | NextSeq | 2X150 | 7908019 |
| **Brassicaceae** | *Matthiola* | *longipetala* | ThermoFisher PureLink RNA Mini Kit | TruSeq | NextSeq | 2X75 | 18984562 | Qiagen DNeasy | TruSeq | NextSeq | 2X150 | 10044883 |
| **Brassicaceae** | *Meniocus* | *linifolius* | ThermoFisher PureLink RNA Mini Kit | TruSeq | NextSeq | 2X75 | 51075472 | Qiagen DNeasy | TruSeq | NextSeq | 2X150 | 9512838 |
| **Brassicaceae** | *Myagrum* | *perfoliatum* | ThermoFisher PureLink RNA Mini Kit | TruSeq | HiSeq | 2X100 | 6922905 | Qiagen DNeasy | TruSeq | NextSeq | 2X150 | 6637717 |
| **Brassicaceae** | *Nasturtium* | *officinale* | ThermoFisher PureLink RNA Mini Kit | TruSeq | NextSeq | 2X75 | 52146124 | Qiagen DNeasy | TruSeq | NextSeq | 2X150 | 8184647 |
| **Brassicaceae** | *Olimoarabidopsis* | *pumila* | ThermoFisher PureLink RNA Mini Kit | TruSeq | NextSeq | 2X75 | 19934340 | *NA* | *NA* | *NA* | *NA* | *NA* |
| **Brassicaceae** | *Physaria* | *acutifolia* | ThermoFisher PureLink RNA Mini Kit | TruSeq | NextSeq | 2X75 | 21947053 | Qiagen DNeasy | TruSeq | NextSeq | 2X150 | 9285650 |
| **Brassicaceae** | *Psychine* | *stylosa* | ThermoFisher PureLink RNA Mini Kit | TruSeq | NextSeq | 2X75 | 17163612 | Qiagen DNeasy | TruSeq | NextSeq | 2X150 | 9336172 |
| **Brassicaceae** | *Rorippa* | *islandica* | ThermoFisher PureLink RNA Mini Kit | TruSeq | HiSeq | 2X100 | 7662263 | Qiagen DNeasy | TruSeq | NextSeq | 2X150 | 7161563 |
| **Brassicaceae** | *Schizopetalum* | *walkeri* | ThermoFisher PureLink RNA Mini Kit | TruSeq | NextSeq | 2X75 | 16524348 | Qiagen DNeasy | TruSeq | NextSeq | 2X150 | 9736577 |
| **Brassicaceae** | *Sinapis* | *alba* | ThermoFisher PureLink RNA Mini Kit | TruSeq | HiSeq | 2X100 | 7602469 | Qiagen DNeasy | TruSeq | NextSeq | 2X150 | 9902350 |
| **Brassicaceae** | *Sisymbrium* | *brassiciforme* | ThermoFisher PureLink RNA Mini Kit | TruSeq | NextSeq | 2X75 | 22582931 | Qiagen DNeasy | TruSeq | NextSeq | 2X150 | 9078512 |
| **Brassicaceae** | *Sisymbrium* | *leucocladum* | ThermoFisher PureLink RNA Mini Kit | TruSeq | NextSeq | 2X75 | 15845037 | Qiagen DNeasy | TruSeq | NextSeq | 2X150 | 10653471 |
| **Brassicaceae** | *Sisymbrium* | *sp* | ThermoFisher PureLink RNA Mini Kit | TruSeq | NextSeq | 2X75 | 16438394 | *NA* | *NA* | *NA* | *NA* | *NA* |
| **Brassicaceae** | *Strepthanthus* | *arizonicus* | ThermoFisher PureLink RNA Mini Kit | TruSeq | NextSeq | 2X75 | 17359223 | Qiagen DNeasy | TruSeq | NextSeq | 2X150 | 10016265 |
| **Brassicaceae** | *Strepthanthus* | *heterophyllus* | ThermoFisher PureLink RNA Mini Kit | TruSeq | NextSeq | 2X75 | 17818482 | Qiagen DNeasy | TruSeq | NextSeq | 2X150 | 10220458 |
| **Brassicaceae** | *Teesdalia* | *nudicaulis* | ThermoFisher PureLink RNA Mini Kit | TruSeq | NextSeq | 2X75 | 59723745 | Qiagen DNeasy | TruSeq | NextSeq | 2X150 | 8579441 |
| **Brassicaceae** | *Thlaspi* | *arvense* | ThermoFisher PureLink RNA Mini Kit | TruSeq | HiSeq | 2X100 | 7651483 | Qiagen DNeasy | TruSeq | NextSeq | 2X150 | 6871390 |
| **Brassicaceae** | *Turritis* | *glabra* | ThermoFisher PureLink RNA Mini Kit | TruSeq | NextSeq | 2X75 | 15406084 | Qiagen DNeasy | TruSeq | NextSeq | 2X150 | 10860792 |
| **Capparaceae** | *Boscia* | *sp* | ThermoFisher PureLink RNA Mini Kit | TruSeq | NextSeq | 2X75 | 17980307 | Qiagen DNeasy | TruSeq | NextSeq | 2X150 | 10870502 |
| **Capparaceae** | *Cadaba* | *natalensis* | ThermoFisher PureLink RNA Mini Kit | TruSeq | NextSeq | 2X75 | 18255177 | Qiagen DNeasy | TruSeq | NextSeq | 2X150 | 9366417 |
| **Capparaceae** | *Capparaceae* | *sp* | ThermoFisher PureLink RNA Mini Kit | TruSeq | NextSeq | 2X75 | 17848365 | Qiagen DNeasy | TruSeq | NextSeq | 2X150 | 9425035 |

| | | | | | | | | | | | | |
|---|---|---|---|---|---|---|---|---|---|---|---|---|
| **Capparaceae** | *Capparis* | *fascicularis* | ThermoFisher PureLink RNA Mini Kit | TruSeq | NextSeq | 2X75 | 18084217 | Qiagen DNeasy | TruSeq | NextSeq | 2X150 | 10378771 |
| **Cariaceae** | *Carica* | *papaya* | ThermoFisher PureLink RNA Mini Kit | TruSeq | NextSeq | 2X75 | 18870089 | Qiagen DNeasy | TruSeq | NextSeq | 2X150 | 13335392 |
| **Cleomaceae** | *Arivela* | *viscosa* | Qiagen RNEasy | TruSeq | HiSeq | 2X100 | 23245927 | Qiagen DNeasy | TruSeq | NextSeq | 2X150 | 9098547 |
| **Cleomaceae** | *Cleomaceae* | *sp* | Qiagen RNEasy | TruSeq | HiSeq | 2X100 | 25851821 | Qiagen DNeasy | TruSeq | NextSeq | 2X150 | 10240576 |
| **Cleomaceae** | *Cleome* | *africana* | Qiagen RNEasy | TruSeq | HiSeq | 2X100 | 21205456 | Qiagen DNeasy | TruSeq | NextSeq | 2X150 | 7267839 |
| **Cleomaceae** | *Cleome* | *amblycocarpa* | Qiagen RNEasy | TruSeq | HiSeq | 2X100 | 21519053 | *NA* | *NA* | *NA* | *NA* | *NA* |
| **Cleomaceae** | *Cleome* | *arabica* | ThermoFisher PureLink RNA Mini Kit | TruSeq | NextSeq | 2X75 | 14426573 | Qiagen DNeasy | TruSeq | NextSeq | 2X150 | 7960574 |
| **Cleomaceae** | *Cleome* | *violacea* | ThermoFisher PureLink RNA Mini Kit | TruSeq | HiSeq | 2X250 | 6720824 | Qiagen DNeasy | TruSeq | NextSeq | 2X150 | 7683480 |
| **Cleomaceae** | *Cleomella* | *serrulata* | Qiagen RNEasy | TruSeq | | | | *NA* | *NA* | *NA* | *NA* | *NA* |
| **Cleomaceae** | *Coalisina* | *angustifolia* | Qiagen RNEasy | TruSeq | HiSeq | 2X100 | 19351183 | Qiagen DNeasy | TruSeq | NextSeq | 2X150 | 8842965 |
| **Cleomaceae** | *Coalisina* | *paradoxa* | Qiagen RNEasy | TruSeq | HiSeq | 2X100 | 21871851 | Qiagen DNeasy | TruSeq | NextSeq | 2X150 | 10046879 |
| **Cleomaceae** | *Gynandropsis* | *gynandra* | Qiagen RNEasy | TruSeq | HiSeq | 2X100 | 19102136 | Qiagen DNeasy | TruSeq | NextSeq | 2X150 | 9236435 |
| **Cleomaceae** | *Melidiscus* | *giganteus* | Qiagen RNEasy | TruSeq | HiSeq | 2X100 | 19954871 | Qiagen DNeasy | TruSeq | NextSeq | 2X150 | 9018300 |
| **Cleomaceae** | *Polanisia* | *dodecandra* | Qiagen RNEasy | TruSeq | HiSeq | 2X100 | 44674939 | Qiagen DNeasy | TruSeq | NextSeq | 2X150 | 8796223 |
| **Cleomaceae** | *Polanisia* | *graveolens* | Qiagen RNEasy | TruSeq | HiSeq | 2X100 | 25106398 | Qiagen DNeasy | TruSeq | NextSeq | 2X150 | 10218176 |
| **Cleomaceae** | *Polanisia* | *sp* | Qiagen RNEasy | TruSeq | HiSeq | 2X100 | 25823291 | Qiagen DNeasy | TruSeq | NextSeq | 2X150 | 8242370 |
| **Cleomaceae** | *Polanisia* | *trachysperma* | Qiagen RNEasy | TruSeq | HiSeq | 2X100 | 38182441 | Qiagen DNeasy | TruSeq | NextSeq | 2X150 | 9082194 |
| **Cleomaceae** | *Sieruela* | *monophylla* | Qiagen RNEasy | TruSeq | HiSeq | 2X100 | 25080383 | *NA* | *NA* | *NA* | *NA* | *NA* |
| **Cleomaceae** | *Tarenaya* | *hassleriana* | Qiagen RNEasy | TruSeq | HiSeq | 2X100 | 19162929 | Qiagen DNeasy | TruSeq | NextSeq | 2X150 | 12209330 |
| **Moringaceae** | *Moringa* | *oleifera* | ThermoFisher PureLink RNA Mini Kit | TruSeq | NextSeq | 2X75 | 57179455 | Qiagen DNeasy | TruSeq | NextSeq | 2X150 | 10239447 |
| **Resadaceae** | *Ochradenus* | *baccatus* | ThermoFisher PureLink RNA Mini Kit | TruSeq | NextSeq | 2X75 | 18868053 | Qiagen DNeasy | TruSeq | NextSeq | 2X150 | 9340353 |
| **Resadaceae** | *Reseda* | *ordorata* | ThermoFisher PureLink RNA Mini Kit | TruSeq | NextSeq | 2X75 | 18754089 | Qiagen DNeasy | TruSeq | NextSeq | 2X150 | 10029969 |

| Species Information | | | DupPipe BIC Scores | | | | FASTKs BIC Scores | | | |
|---|---|---|---|---|---|---|---|---|---|---|
| **Family** | **Genus** | **Species** | **1 peak** | **2 peaks** | **3 peaks** | **4 peaks** | **1 peak** | **2 peaks** | **3 peaks** | **4 peaks** |
| **Bataceae** | *Batis* | *maritima* | -6273.05 | -4970.69 | -4612.552 | -4515.273 | -29659.81 | -29227.93 | -28742.21 | -28355.45 |
| **Brassicaceae** | *Aethionema* | *arabicum* | -17888.73 | -15930.41 | -15712.96 | -15489.97 | -72664.46 | -71361.21 | -70739.96 | -70405.8 |
| **Brassicaceae** | *Barbarea* | *bracteosa* | -20416.34 | -18389.58 | -18047.26 | -17929.21 | -55743.7 | -54258.87 | -53867.64 | -53868.36 |
| **Brassicaceae** | *Cakile* | *maritima* | -49327.51 | -48717.18 | -48587.63 | -48384.6 | -140948 | -139956.7 | -139670.2 | -139216.5 |
| **Brassicaceae** | *Calepina* | *irregularis* | -34320.8 | -30254.13 | -29871.31 | -29871.35 | -67744.52 | -64841.66 | -63941.85 | -63706.6 |
| **Brassicaceae** | *Capsella* | *bursa-pastoris* | -9387.535 | -8388.332 | -8312.341 | -8217.367 | -51871.77 | -50359.75 | -49734.31 | -49607.75 |
| **Brassicaceae** | *Cardamine* | *hirsuta* | -18046.43 | -15950.79 | -15655.47 | -15519.12 | -27724.56 | -27082.3 | -26877.96 | -26843.1 |
| **Brassicaceae** | *Caulanthus* | *amplexicaulis* | -72240.47 | -66365.55 | -65466.22 | -65312.87 | -296250.7 | -288865.5 | -286690 | -286481.9 |
| **Brassicaceae** | *Chorispora* | *tenella* | -14681.18 | -13104.49 | -12870.93 | -12797.43 | -27157.47 | -26754.78 | -26675.19 | -26592.95 |
| **Brassicaceae** | *Cochlearia* | *officinalis* | -26835.57 | -25203.31 | -24977.02 | -24724.24 | -45997.75 | -45297.29 | -44895.66 | -44896.44 |
| **Brassicaceae** | *Crambe* | *hispanica* | -53282.57 | -52870.1 | -52509.33 | -52201.88 | -162684.3 | -161909.2 | -161518.4 | -160916.1 |
| **Brassicaceae** | *Descurainia* | *sophioides* | -20134.61 | -17945.48 | -17617.56 | -17430.33 | -78315.54 | -76328.78 | -74915.19 | -74615.81 |
| **Brassicaceae** | *Descurainia* | *pinnata* | -32177.51 | -29224.79 | -28697.38 | -28298.75 | -139705.5 | -136187.3 | -134855.7 | -134853.8 |
| **Brassicaceae** | *Diptycocarpus* | *strictus* | -20176.44 | -18385.49 | -18151.68 | -18005.14 | -51593.24 | -49629.33 | -49558.3 | -49397.94 |
| **Brassicaceae** | *Eruca* | *vesicaria* | -57302.99 | -56279.26 | -56171.14 | -55940.44 | -180363.8 | -178936.7 | -178670.7 | -178002.3 |
| **Brassicaceae** | *Erysimum* | *cheiranthoides* | -17035.26 | -15470.34 | -15201.41 | -15048.28 | -27870.51 | -27251.97 | -27169.5 | -27167.07 |
| **Brassicaceae** | *Euclidium* | *syriacum* | -7330.347 | -6664.564 | -6569.126 | -6486.873 | -75457 | -74132.51 | -74132.3 | -73955.92 |
| **Brassicaceae** | *Farsetia* | *aegyptia* | -87046.73 | -78626.97 | -77752.59 | -77752.63 | -27584.64 | -26808.89 | -26648.29 | -26461.84 |
| **Brassicaceae** | *Goldbachia* | *laevigata* | -46724.37 | -40469.43 | -39908.4 | -39824.05 | -126148.9 | -120596.6 | -119013.2 | -118631.9 |
| **Brassicaceae** | *Guillenia* | *lasiophylla* | -59735.11 | -53526.87 | -52969.1 | -52857.97 | -134706.6 | -128952.9 | -128027.7 | -127946 |
| **Brassicaceae** | *Hesperis* | *matronalis* | -8981.003 | -8402.243 | -8296.301 | -8279.953 | -116152.4 | -114019.3 | -113171.9 | -113171.2 |
| **Brassicaceae** | *Hirschfeldia* | *incana* | -57478.96 | -56124.2 | -56066.99 | -55528.43 | -129935.9 | -129659.6 | -129543 | -128886.2 |
| **Brassicaceae** | *Iberis* | *amara* | -37998.83 | -36103.28 | -35797.37 | -35513.78 | -104300.9 | -102190.1 | -101688.1 | -101659.2 |
| **Brassicaceae** | *Isatis* | *lusitanica* | -16637.79 | -14632.94 | -14324.95 | -14221.36 | -27652.87 | -26943.13 | -26871.77 | -26692.73 |

| | | | | | | | | | | |
|---|---|---|---|---|---|---|---|---|---|---|
| **Brassicaceae** | *Isatis* | *tinctoria* | -32807.62 | -28987.6 | -28672.49 | -28416.86 | -64911.3 | -62615.86 | -61966.98 | -61638.4 |
| **Brassicaceae** | *Lepidium* | *ruderale* | -51478.63 | -43198.8 | -42721.1 | -42559.56 | -125387.6 | -118218.6 | -117136.5 | -116627.7 |
| **Brassicaceae** | *Lepidium* | *sativum* | -63920.94 | -57354.73 | -56523.84 | -56506.45 | -235772.5 | -226791.3 | -224613.5 | -224338.9 |
| **Brassicaceae** | *Lobularia* | *maritima* | -22482.68 | -20460.54 | -20234 | -20233.39 | -35962.92 | -35163.19 | -34845.46 | -34845.36 |
| **Brassicaceae** | *Lunaria* | *annua* | -49086.82 | -48726.38 | -48391.74 | -48324.15 | -156452.4 | -156020.2 | -155609.5 | -155183.2 |
| **Brassicaceae** | *Malcomia* | *maritima* | -31503.91 | -28447.88 | -28044.13 | -27721.6 | -104663.4 | -100716.8 | -100089.1 | -99712.58 |
| **Brassicaceae** | *Matthiola* | *longipetala* | -18787.75 | -16630.62 | -16436.26 | -16259.59 | -28755.13 | -28074.03 | -27838.22 | -27791.81 |
| **Brassicaceae** | *Meniocus* | *linifolius* | -93350.47 | -79319.88 | -78028.4 | -77924.67 | -293255.5 | -277515.4 | -274443.6 | -273907.6 |
| **Brassicaceae** | *Myagrum* | *perfoliatum* | -18501.89 | -16611.93 | -16246.46 | -16074.25 | -46087.16 | -44153.4 | -43545.81 | -43274.85 |
| **Brassicaceae** | *Nasturtium* | *officinale* | -60471.95 | -52138.19 | -51393.83 | -51165.92 | -223823.9 | -213019.7 | -211035.3 | -210681.9 |
| **Brassicaceae** | *Olimoarabidopsis* | *pumila* | -39183.53 | -33871.53 | -33459.93 | -33419.88 | -66715.53 | -63165.63 | -62249.17 | -62076.87 |
| **Brassicaceae** | *Physaria* | *acutifolia* | -41432.66 | -40148.94 | -40051.83 | -39542.31 | -100664.9 | -100252.2 | -100064.3 | -99552.98 |
| **Brassicaceae** | *Psychine* | *stylosa* | -42292.4 | -41469.83 | -41414.98 | -41051.42 | -109069.2 | -108968.6 | -108891.8 | -108786.9 |
| **Brassicaceae** | *Rorippa* | *islandica* | -22644.09 | -20601.65 | -20221.54 | -20067.94 | -53588.67 | -51903.39 | -51623.81 | -51592.38 |
| **Brassicaceae** | *Schizopetalum* | *walkeri* | -42672.93 | -41045.12 | -40688.12 | -40594.02 | -134672.9 | -133861.7 | -132864.5 | -132864.6 |
| **Brassicaceae** | *Sinapis* | *alba* | -58678.87 | -58381.11 | -57891.62 | -57599.67 | -214304.1 | -213673.6 | -213076.5 | -212988.6 |
| **Brassicaceae** | *Sisymbrium* | *brassiciforme* | -56912.82 | -51040.69 | -50499.11 | -50441.32 | -106873.4 | -106872.7 | -106631 | -106195.7 |
| **Brassicaceae** | *Sisymbrium* | *leucocladum* | -18171.84 | -16346.14 | -16073.08 | -15879.77 | -29525.35 | -28937.77 | -28532.62 | -28429.9 |
| **Brassicaceae** | *Sisymbrium* | *sp* | -45927.74 | -44982.75 | -44902.77 | -44505.49 | -48002.97 | -45832.08 | -45213.05 | -44905.1 |
| **Brassicaceae** | *Brassicaceae* | *sp* | -92159.93 | -85720.01 | -84367.26 | -84131.4 | -397328.4 | -390193.2 | -386693.6 | -385884.6 |
| **Brassicaceae** | *Strepthanthus* | *arizonicus* | -44070.05 | -41193.34 | -40805.58 | -40760.63 | -92667.5 | -89997.98 | -89422.72 | -89310.5 |
| **Brassicaceae** | *Strepthanthus* | *heterophyllus* | -47146.15 | -43765.39 | -43355.83 | -43327.92 | -98277.5 | -95744.79 | -94996.69 | -94933.27 |
| **Brassicaceae** | *Teesdalia* | *nudicaulis* | -70434.89 | -62882.51 | -41633.38 | -61916.37 | -210073 | -202074 | -199908.6 | -199724.9 |
| **Brassicaceae** | *Thlaspi* | *arvense* | -21362.18 | -19322.8 | -18939.83 | -18772.49 | -54264.15 | -53094.77 | -53083.27 | -52492.75 |
| **Brassicaceae** | *Turritis* | *glabra* | -49465.03 | -42282.28 | -41683.33 | -41633.38 | -92297.93 | -86449.15 | -85442.42 | -85345.57 |
| **Capparaceae** | *Boscia* | *sp* | -15462.33 | -13656.09 | -12905.13 | -12735.44 | -57128.34 | -55238.59 | -54170.86 | -54008.21 |
| **Capparaceae** | *Cadaba* | *natalensis* | -10654.85 | -8824.122 | -8174.672 | -8072.744 | -47515.81 | -45670.41 | -45094.29 | -44846.13 |
| **Capparaceae** | *Capparis* | *fascicularis* | -13851.04 | -12299.53 | -11690.69 | -11549.21 | -43810.82 | -42563.49 | -41808.2 | -41170.72 |

| | | | | | | | | | | |
|---|---|---|---|---|---|---|---|---|---|---|
| **Capparaceae** | *Capparis* | *tomentosa* | -14017.43 | -12372.35 | -11778.9 | -11641.74 | -55875.39 | -54956.17 | -54456.94 | -54060.24 |
| **Cariaceae** | *Carica* | *papaya* | -12964.8 | -10810.21 | -10361.01 | -10318.83 | -37739.28 | -36667.15 | -35950.16 | -35652.17 |
| **Cleomaceae** | *Cleomaceae* | *sp* | -32790.84 | -31907.16 | -30767.01 | -30457.21 | -121041.3 | -120137.4 | -119236.3 | -118223 |
| **Cleomaceae** | *Cleome* | *africana* | -12033.86 | -10203.68 | -9700.341 | -9570.846 | -54351.44 | -53335.11 | -53115.26 | -52575.36 |
| **Cleomaceae** | *Cleome* | *amblycocarpa* | -10307.71 | -8534.686 | -8097.695 | -8023.899 | -39832.76 | -38657.31 | -37793.17 | -37665.87 |
| **Cleomaceae** | *Cleome* | *arabica* | -7773.53 | -6276.361 | -5974.505 | -5878.58 | -15333.58 | -14779.72 | -14518.46 | -14386.6 |
| **Cleomaceae** | *Sieruela* | *monophylla* | -34499.78 | -33835.05 | -32662.68 | -32407.19 | -125206.2 | -124041.9 | -124342.1 | -123520.9 |
| **Cleomaceae** | *Cleome* | *violacea* | -14478.17 | -12665.1 | -12143.09 | -12031.81 | -49680.57 | -48555.81 | -48077.79 | -48054.83 |
| **Cleomaceae** | *Melidiscus* | *giganteus* | -39082.77 | -38144.81 | -37158.06 | -36845.59 | -208644 | -207264.7 | -206914.5 | -204958.6 |
| **Cleomaceae** | *Arivela* | *viscosa* | -12125.13 | -10601.82 | -10094.69 | -9995.74 | -46372.67 | -44884.62 | -44811.79 | -44461.78 |
| **Cleomaceae** | *Cleomella* | *lutea* | -11321.78 | -9636.747 | -9186.818 | -9072.467 | -46405.07 | -45258.65 | -44304.96 | -43969.71 |
| **Cleomaceae** | *Cleomella* | *serrulata* | -35545.31 | -29633.1 | -28966.29 | -28500.9 | -104059.2 | -98769.64 | -97286.63 | -96448.16 |
| **Cleomaceae** | *Coalisina* | *angustifolia* | -32947.1 | -32573.87 | -31485.88 | -31275.91 | -70695.18 | -70690.09 | -70254.17 | -69824.07 |
| **Cleomaceae** | *Coalisina* | *paradoxa* | -45113.69 | -43013.85 | -42454.3 | -42123.13 | -232073.7 | -230283.9 | -228074.1 | -226942.3 |
| **Cleomaceae** | *Gynandropsis* | *gynandra* | -26445.88 | -25136.17 | -24368.74 | -24136.14 | -105952.6 | -104635 | -103678.3 | -103482.8 |
| **Cleomaceae** | *Polanisia* | *dodecandra* | -15250.73 | -13157.53 | -12519.87 | -12362.36 | -62510.34 | -61483.52 | -60632.61 | -60304.93 |
| **Cleomaceae** | *Polanisia* | *graveolens* | -11921.91 | -9855.295 | -9345.755 | -9212.286 | -33712.79 | -32733.65 | -32274.19 | -31957.47 |
| **Cleomaceae** | *Polanisia* | *trachysperma* | -13878.81 | -11855.59 | -11252.52 | -11116.56 | -56577.94 | -55015.23 | -54726.4 | -54212.57 |
| **Cleomaceae** | *Tarenaya* | *hassleriana* | -34991.25 | -34328.86 | -33198.46 | -33171.58 | -113691.1 | -113078.4 | -112644.8 | -112107.3 |
| **Moringaceae** | *Moringa* | *oleifera* | -8975.359 | -6567.789 | -6563.584 | -6244.213 | -19414.45 | -18830.46 | -18389.46 | -18135.18 |
| **Resadaceae** | *Ochradenus* | *baccatus* | -30000.1 | -28976.74 | -28050.13 | -27748.79 | -65666.52 | -64702.93 | -64643.43 | -63693.03 |
| **Resadaceae** | *Reseda* | *ordorata* | -29667.79 | -28450.24 | -27615.78 | -27382.23 | -68356.46 | -67711.71 | -67638.78 | -67120.38 |
